# DREADD-mediated amygdala activation is sufficient to induce anxiety-like responses in young nonhuman primates

**DOI:** 10.1101/2023.06.06.543911

**Authors:** Sascha A.L. Mueller, Jonathan A. Oler, Patrick H. Roseboom, Nakul Aggarwal, Margaux M. Kenwood, Marissa K. Riedel, Victoria R. Elam, Miles E. Olsen, Alexandra H. DiFilippo, Bradley T. Christian, Xing Hu, Adriana Galvan, Matthew A. Boehm, Michael Michaelides, Ned H. Kalin

## Abstract

Anxiety disorders are among the most prevalent psychiatric disorders, with symptoms often beginning early in life. To model the pathophysiology of human pathological anxiety, we utilized Designer Receptors Exclusively Activated by Designer Drugs (DREADDs) in a nonhuman primate model of anxious temperament to selectively increase neuronal activity of the amygdala. Subjects included 10 young rhesus macaques; 5 received bilateral infusions of AAV5-hSyn-HA-hM3Dq into the dorsal amygdala, and 5 served as controls. Subjects underwent behavioral testing in the human intruder paradigm following clozapine or vehicle administration, prior to and following surgery. Behavioral results indicated that clozapine treatment post-surgery increased freezing across different threat-related contexts in hM3Dq subjects. This effect was again observed approximately 1.9 years following surgery, indicating the long-term functional capacity of DREADD-induced neuronal activation. [^11^C]deschloroclozapine PET imaging demonstrated amygdala hM3Dq-HA specific binding, and immunohistochemistry revealed that hM3Dq-HA expression was most prominent in basolateral nuclei. Electron microscopy confirmed expression was predominantly on neuronal membranes. Together, these data demonstrate that activation of primate amygdala neurons is sufficient to induce increased anxiety-related behaviors, which could serve as a model to investigate pathological anxiety in humans.

## Introduction

Fear and anxiety have evolved as important adaptive responses to promote the survival of an organism when confronted with environmental threats. Individual differences in the expression of anxiety-related behaviors can be observed early in life, and high levels of trait anxiety, or extreme behavioral inhibition, are a prominent risk factor for the development of anxiety disorders (ADs) (Clauss & Blackford, 2012; Hirshfeld-Becker et al., 2008). ADs are characterized by extreme subjective distress and avoidance behaviors that persist even in non-threatening situations and contexts (Dymond et al., 2015). There is considerable evidence implicating the dysregulation of specific neural circuits that mediate threat-related defensive behaviors in the pathophysiology of ADs (Daviu et al., 2019; Duval et al., 2015; McTeague et al., 2020). In this regard, we developed a nonhuman primate (NHP) model of anxious temperament to further investigate mechanisms underlying the neural alterations and pathological behaviors that are associated with ADs in humans (Fox et al., 2008). The NHP model is ideally suited for this purpose, due to shared similarities to humans in behavior as well as brain structure and function (Kalin & Shelton, 2003).

In both humans and NHPs, individual differences in the expression of trait-like anxiety-related behaviors emerge early in life and exist along a continuum of severity (Fox, Oler, Shackman, et al., 2015). Young rhesus monkeys on the high end of this continuum are similar to children with high levels of behavioral inhibition, and we have demonstrated that extreme trait-like anxiety in young monkeys can serve as a translational model for the childhood risk to develop anxiety and other comorbid disorders (Fox & Kalin, 2014). Additionally, using this model we found that increased metabolism in the dorsal amygdala (which includes the central nucleus [Ce], as well as dorsal aspects of the basal and accessory basal nuclei) and other limbic, prefrontal, and brainstem regions, is associated with extreme anxious temperament (Fox et al., 2008; Fox, Oler, Shackman, et al., 2015). This is consistent with human fMRI studies demonstrating increased amygdala activity in individuals with ADs (Chavanne & Robinson, 2021; Engel et al., 2009; Etkin et al., 2004; Etkin & Wager, 2007; Fredrikson & Faria, 2013; Holzschneider & Mulert, 2011; Ipser et al., 2013).

Previous studies using lesion (Kalin et al., 2004; Machado & Bachevalier, 2008) and chemogenetic strategies (Raper et al., 2019; Roseboom et al., 2021) to decrease amygdala activity in young NHPs have demonstrated a reduction in anxious behaviors, thereby mechanistically implicating the amygdala as having a key role in mediating adaptive anxiety-related responses. The chemogenetic studies have been particularly informative, as they allowed for the reversible modulation of amygdala neurons. To further understand the role of the NHP amygdala in mediating pathological and maladaptive anxiety, here we performed experiments with hM3Dq DREADDs (Designer Receptors Exclusively Activated by Designer Drugs) to assess the effects of acute amygdala activation on anxiety-related behaviors expressed in response to the human intruder paradigm (HIP). The HIP consists of three different fear-related contexts, including: Alone, No-Eye-Contact (NEC), and Stare, which elicit different defensive responses (Kalin & Shelton, 1989). During Alone, the typical responses are increased cooing and locomotion, and these function to increase the likelihood of reuniting with a conspecific. The NEC condition, in which a human presents his/her profile to the monkey, is perceived as a potential threat and typically results in reduced locomotion, increased freezing behavior, and a reduction in vocalizations. The Stare condition, in which the human intruder stares directly at the monkey with a neutral face, is perceived as a direct threat and elicits a combination of aggressive and submissive gestures from the monkey. Behavioral testing was performed at two timepoints, approximately two years apart, to demonstrate the replicability of the findings, as well as to assess the extent to which DREADD technology remained functional over a prolonged time period. Finally, we used positron emission tomography (PET) and the [^11^C]deschloroclozapine ([^11^C]DCZ) radiotracer, along with postmortem immunohistochemical and electron microscopy studies, to verify and characterize DREADD expression.

The overall aim of these studies is to provide evidence in NHPs for a mechanistic role of increased amygdala activity in mediating the pathophysiology underlying pathological anxiety. Because this NHP model is highly translational, these studies have the potential to provide unique insights into treatment strategies for pathological anxiety in humans.

## Methods

### Subject details

The present study was performed using 10 female rhesus monkeys (*Macaca mulatta)*, 5 of which were randomly selected to undergo intraoperative MRI (iMRI) surgery and become hM3Dq subjects, while the other 5 served as unoperated controls. Subjects were housed in pairs, with each pair consisting of an hM3Dq subject and an unoperated control subject. Monkeys were housed and cared for at the Harlow Center for Biological Psychology on a 12-hour light/dark cycle, in a temperature- and humidity-controlled vivarium. Procedures were performed using protocols that were approved by the University of Wisconsin Institutional Animal Care and Use Committee. At the time of iMRI surgery, hM3Dq subjects and their cagemate controls were, on average, 1.73 (± 0.09) years old, an age range that falls within preadolescence. See Supplemental Table 1 for information regarding which experiments were performed with a given subject.

### Experiment 1: Behavioral testing with clozapine and vehicle

The timeline for behavioral testing is shown in Figure 1. To maximize the potential for observing the increase in freezing that was hypothesized to result from hM3Dq activation, we attempted to select animals that displayed low-mid levels of freezing during pre-testing. Of the 64 animals that were initially screened (30 males/34 females, mean age at time of screening 1.42 ± 0.19 years), 16 were then chosen to be tested for responsiveness to clozapine. The pre-surgical, or “baseline,” testing period consisted of 10 minutes of Alone, followed by 10 minutes of NEC. Using the same individualized dosing strategy as in Roseboom et al. (2021), baseline testing consisted of three instances of behavioral assessment, in which 0.03 mg/kg clozapine, 0.1 mg/kg clozapine, or vehicle were administered (IM) 30 minutes prior to testing. The solutions used for the 0.03 mg/ kg and 0.1 mg/kg doses were prepared at a concentration of 0.06 mg/ mL or 0.2 mg/mL, respectively, by dissolving the clozapine (Sigma-Aldrich, St. Louis, MO) in DMSO and then diluting in 0.9% saline to a final DMSO concentration of 25%. The drug was administered as bilateral intramuscular (IM) injections at a volume of 0.5 mL/kg. Vehicle injected animals received 0.5 ml/kg of 25% DMSO in 0.9% saline. The dose of clozapine chosen for each pair of subjects to be used during post-surgical testing was determined based on freezing behavior observed during the pre-testing period. The highest dose of clozapine that did not appear to substantially affect freezing during the pre-testing NEC context was selected for each pair of animals. 0.1 mg/kg clozapine was selected for one pair of subjects (consisting of subjects E1 and C1), while 0.03 mg/kg clozapine was selected for the other four pairs. Neuroendocrine effects of clozapine at both doses were also assessed using blood collected from each subject (refer to Supplemental Materials and Supplemental Figure 1 for details). Blood collection was performed at approximately the same time of day, immediately following the end of each baseline behavioral testing period (range 08:16 AM – 10:49 AM).

**Figure 1.**
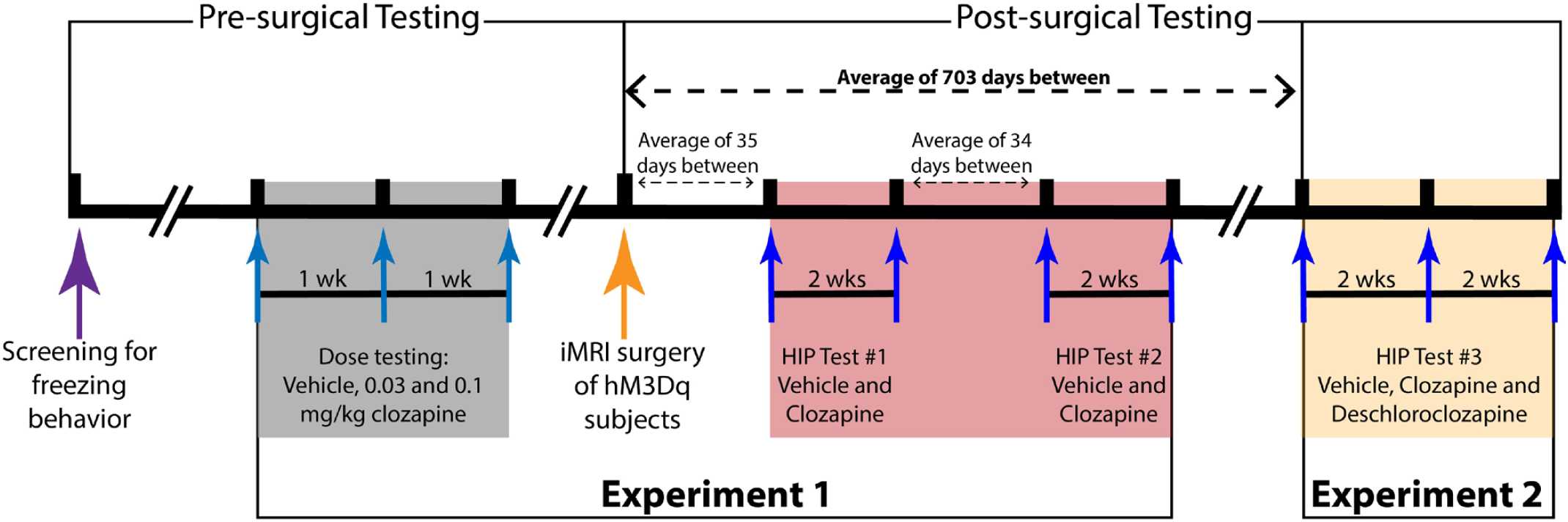
Timeline of behavioral testing. 10 subjects were included in Experiment 1 (n = 5 hM3Dq subjects and 5 unoperated cagemate controls). 8 of these subjects were included in Experiment 2 (n = 4 hM3Dq subjects and 4 unoperated cagemate controls.)

Post-surgical testing began approximately 35 (± 3) days after iMRI surgery. Behavioral testing during the post-surgical period consisted of the following contexts, listed in order of occurrence: 10 min of Alone, 10 min of NEC, 5 min of Alone, 10 min of Stare, and 15 min of Alone. At the end of testing, blood collection was performed at approximately the same time of day (range 08:55 AM – 09:28 AM). Clozapine and vehicle administration were randomized, with at least a period of two weeks between each instance of testing. Subjects underwent a second round of HIP testing approximately 34 (± 2) days after the completion of the first round of HIP testing (or 83 ± 2 days after iMRI surgery), and administration of clozapine and vehicle was counterbalanced, again with a period of at least two weeks between each instance of testing. Of note, the behavioral data from the first round of HIP was lost for one subject pair (E1 and C1), and so this pair of subjects underwent an additional “third” round of HIP 114 days after completion of their second round of HIP testing (or 212 days after iMRI surgery of subject E1).

### Experiment 2: Behavioral testing with clozapine, deschloroclozapine, and vehicle following long-term DREADD expression

4 pairs of subjects (E2-5 and C2-5; see Supplemental Table 1) underwent further HIP testing on vehicle, clozapine, and deschloroclozapine (DCZ), starting approximately 703 (± 38) days after iMRI surgery of the hM3Dq subjects. Because clozapine and DCZ have similar selectivity and affinity for the hM3Dq receptor ((Nagai et al., 2020; Yan et al., 2021)), the dose of DCZ utilized here was the same as the dose of clozapine chosen for post-surgical testing in Experiment 1, which for all subjects in this experiment (n = 8) was 0.03 mg/kg. DCZ (MedChemExpress, Monmouth Junction, NJ) was prepared by dissolving in dimethyl sulfoxide (DMSO) and then diluting with 0.9% saline to a final concentration of 0.06 mg/ml in 25% DMSO. As previously described for Experiment 1, the drug was administered IM in a volume of 0.5 ml/kg. The vehicle solution was 25% DMSO in 0.9% saline and was also injected in a volume of 0.5 ml/kg. Each testing period again consisted of 10 min of Alone, 10 min of NEC, 5 min of Alone, 10 min of Stare, and 15 min of Alone. At the end of testing, blood collection was performed at approximately the same time of day (range 09:06 AM – 09:25 AM). The administration of vehicle or DCZ was randomized in the first two instances of testing, and clozapine was used for all subjects in the third instance, with a period of two weeks separating each test. Drug or vehicle administration was performed 30 minutes prior to testing by means of bilateral intramuscular injections.

It should be noted that in the second 5-minute epoch of NEC (“N2”), testing was shortened by 55.23 seconds for subject E2, and in the third 5-minute epoch of Alone following Stare (“A6”), testing was shortened by 3.18 seconds for subject C5. Therefore, data was extrapolated for these two subjects during those specific contexts (N2 for subject E2, and A6 for subject C5), such that the duration of freezing or locomotion observed was divided by the total number of seconds of data collection and then multiplied by 300 (to represent 5 full minutes of each epoch).

### Statistical Analysis of Behavior

Linear mixed-effects (LME) models were implemented for all behavioral analyses, allowing for precise and unbiased effect estimates by accounting for repeated within-subject measures (Brauer & Curtin, 2018). All LME modeling was performed using the lme4 package in RStudio (ver. 2022.02.1) (Bates et al., 2015). The primary dependent variables of interest included freezing, locomotion, and coo-vocalizations. Additionally, bark-vocalizations and experimenter-directed hostility were examined specifically during the Stare condition. Freezing was defined as a lack of movement for at least 3 seconds, characterized by no vocalizations and no movement other than isolated movements of the head, or slight body movements used to maintain posture. Locomotion was defined as ambulation of one or more full steps at any speed. Experimenter hostility was defined as any hostile behavior directed at the human intruder, e.g., barking, head bobbing, ear flapping, etc. Bark-vocalizations were defined as vocalization made by forcing air through vocal cords from the abdomen, producing a short, rasping low frequency sound. Coo-vocalizations were defined as vocalization made by rounding and pursing the lips, with an initial increase and subsequent decrease in frequency and intensity. To normalize the data, duration behaviors (freezing, locomotion, and experimenter-directed hostility) were log-transformed (log(seconds) + 1), and frequency behaviors (coo and bark vocalizations) were square-root transformed. To account for potential individual differences in response to drug, and because different doses of drug were used for different pairs of animals, data were residualized for blood plasma concentrations of drug (clozapine, or DCZ) using IBM SPSS Statistics v. 28 (IBM, Armonk, NY, USA) prior to statistical analysis. Plasma concentrations for data following vehicle administration were set to zero. Prism v. 9.3.1 (GraphPad Software, San Diego, CA) was used for the preparation of graphical representations.

#### Experiment 1

Pre-surgical baseline data collected for vehicle and the dose of clozapine selected for post-surgical testing were included in the analyses. Independent variables examined with LME analyses were Group (control versus hM3Dq), Treatment (vehicle versus clozapine), Pre/Post (pre-surgical versus post-surgical testing), and Condition (Alone versus NEC.) LME models tested the 4-way interaction of these variables, as well as all lower order effects. Planned comparisons for the significant 3-way interaction of interest (Group x Treatment x Pre/Post) were conducted to examine differences between vehicle and clozapine in each Group at each timepoint. Bonferroni correction was implemented to account for multiple comparisons (α = 0.05/8; p = 0.00625, corrected). This correction was based upon the number of dependent variables found to have the significant 3-way interaction (2: freezing and locomotion) and the number of t-tests performed comparing vehicle to clozapine (4: control Pre, hM3Dq Pre, control Post, hM3Dq Post).

Because the Stare condition was only included during the post-surgical testing period, behaviors collected during this context were analyzed separately, and the independent variables examined with LME analyses were Group and Treatment. Planned comparisons for the significant interaction of interest (Group x Treatment) were conducted to examine differences between vehicle and clozapine in each Group. Bonferroni correction was implemented to account for multiple comparisons (α = 0.05/4; p = 0.0125, corrected), again based upon the number of dependent variables examined (2: freezing and locomotion) and the number of t-tests performed comparing vehicle to clozapine (2: control Post, hM3Dq Post).

#### Experiment 2

Because Experiment 2 was conducted approximately 2 years after presurgical baseline testing, and because baseline testing did not include DCZ, only post-surgical data following long-term DREADD expression was utilized in these analyses. The independent variables examined with the LME analyses included Group (control versus hM3Dq), Treatment (vehicle versus clozapine, or vehicle versus DCZ), and Condition (Alone versus NEC versus Stare). For this experiment, two separate analyses were conducted: one comparing clozapine to vehicle data, and another comparing DCZ to vehicle data. The vehicle data was the same in both analyses. Planned comparisons for either the significant 2-way (Group x Treatment) or 3-way (Group x Treatment x Condition) interactions of interest were conducted to examine differences between vehicle and drug within each Group.

Bonferroni correction was implemented to account for multiple comparisons related to the 2-way interaction of interest (α = 0.05/8; p = 0.00625.) This correction was based upon the number of dependent variables examined (4: freezing, locomotion, and coo-vocalizations following clozapine administration, and freezing following DCZ administration) and the number of t-tests performed comparing vehicle to clozapine (2: control, hM3Dq).

Bonferroni correction was also implemented to account for multiple comparisons related to the 3-way interaction of interest (α = 0.05/6; p = 0.00833.) Again this correction was based upon the number of dependent variables examined (1: locomotion following DCZ administration) and the number of t-tests performed comparing vehicle to clozapine (6: control Alone, control NEC, control Stare, hM3Dq Alone, hM3Dq NEC, hM3Dq Stare).

For Stare-specific behaviors the independent variables included in the LME analyses were Group and Treatment. Bonferroni correction was implemented to account for multiple comparisons (α = 0.05/4; p = 0.0125, corrected), again based upon the number of dependent variables examined (2: bark-vocalizations following both clozapine and DCZ administration) and the number of t-tests performed (2: control, hM3Dq).

### Assessing plasma levels of clozapine, deschloroclozapine, and related metabolites

Plasma levels of clozapine (CLZ), deschloroclozapine (DCZ), norclozapine (NOR), clozapine-N-oxide (CNO), and compound 21 (C21) were processed using methanol protein precipitation and analyzed with HPLC coupled mass spectroscopy as in methods previously described (Bonaventura et al., 2019; Roseboom et al., 2021). A Nexera XR HPLC (Shimadzu, Kyoto, Japan) coupled with a QTRAP 6500+ (SCIEX, Redwood City, CA, USA) was used to acquire multiple reaction monitoring (MRM) data analyzed with Analyst 1.6 (SCIEX). Separation of CLZ, DCZ, NOR, CNO, and C21 was accomplished using a C18 Security guard cartridge (4.6 × 4 mm) and Zorbax Extend-C18 column (4.6 × 75 mm, 3.5 μm, Agilent 766953-902) at 40 °C (mobile phase A = H_2_O + 0.1% formic acid, mobile phase B = MeOH + 0.1% formic acid). The following MRM ion transitions were used for each compound: CLZ (327.30 → 270.10), DCZ (292.96 → 236.10), NOR (312.70 → 227.10), CNO (343.80 → 191.90), C21 (278.97 → 193.00) and internal standard JHU37160 (359.10 → 244.80). Standard calibration curves for each compound were prepared using a 0.5 serial dilution (CLZ and DCZ range 50 ng/ml – 0.0015 ng/ml; NOR, CNO and C21 range 10 ng/ml – 0.0003ng/ml).

### Surgical procedures and viral vector delivery

#### Trajectory guide base placement and intraoperative MRI

The placement of the trajectory guide bases followed published methods (Emborg et al., 2014; Emborg et al., 2010) that were modified to target the dorsal amygdala following procedures described in supplemental methods section of two previous publications (Fox et al., 2019; Kalin et al., 2016). One infusion of 24 µl was performed per hemisphere, with the intent of covering as much of the dorsal amygdala as possible, which includes the Ce and dorsal aspects of the basal and accessory basal nuclei. The infusion was performed in increments, such that a small portion of the 24 µl volume was infused, iMRI scans were collected to visualize the placement of the infusion, and then injector depth was adjusted if necessary; these steps were repeated until a total of 24 µl of the viral vector was infused.

Before the procedure, the animals were anesthetized with ketamine (up to 20 mg/kg, intramuscular (IM)), prepared for surgery, and then placed in an MRI compatible-stereotaxic frame. The animals were intubated and received isoflurane anesthesia (1 – 2%, intratracheal) with 1-1.5% O_2_ administered during induction. Atropine sulfate (0.01 – 0.4 mg/kg, IM) was administered to depress salivary secretion, and buprenorphine (0.01 – 0.03 mg/kg, IM, repeated every 6 – 12 hours) was given for analgesia. To maintain fluids and electrolytes, Plasmalyte (up to 10 mg/kg/hr, intravenous (IV)) was administered. Cefazolin (20 – 25 mg/kg, IM or IV) was administered as a prophylactic antibiotic one day prior to the surgery. Cefazolin was also administered immediately prior to surgery, and then every 6 hours while under anesthesia. All drugs and treatments were given in consultation with veterinary staff. Vital signs (heart rate, respiration, oxygen saturation, non-invasive blood pressure, and end tidal CO_2_) were continuously monitored. Body temperature was monitored during the surgical and MRI procedure and maintained in the MRI by wrapping the animals for warmth while incorporating a hot water heating blanket within the wrap.

#### Cannula trajectory planning and insertion

Cannula trajectory planning was carried out in the MRI suite under anesthesia as previously described (Fox et al., 2019). The infusion line was primed with a loading line solution that was identical to the virus suspension solution [Corning phosphate-buffered saline without Ca^2+^ or Mg^2+^ with 350 mM NaCl (Gibco, ThermoFisher Scientific, Waltham, MA) containing 0.001% F68 surfactant (Gibco, ThermoFisher Scientific)]. The cannula was loaded with the DREADD viral vector which was mixed with the MR visible marker Gadobenate dimeglumine (Gd, MultiHance; Bracco Diagnostics, Monroe Township, NJ). A 100 mm length cannula (Engineering Resources Group) with a fused silica-end port cannula (OD = 0.67 mm and ID = 0.45 mm) was utilized for viral vector infusion in all five hM3Dq-HA subjects. After pressure in the infusion line was checked and stabilized, the cannula was introduced into the brain by advancing the remote introducer at a rate of approximately 10 – 15 mm/minute. The cannula was advanced two-thirds of the measured depth toward the chosen target, at which point a 3D T1W MRI was performed to confirm the correct trajectory and to aid in estimating the remaining distance between the cannula tip and the target. The cannula was then advanced to the target position, and when the pressure measured by the infusion pump controller was stable, the infusion was initiated.

The infusate consisted of AAV5-hSyn-HA-hM3Dq suspended in a solution identical to the loading line solution described above. *In vivo* MRI visualization of the infusion was made possible with Gd, which was mixed with the viral vector at a final concentration of 0.66 mM. Each hM3Dq subject received a total volume of 24 μl per hemisphere, infused at a rate of 1 μl/min. After infusion into each hemisphere was complete, a 3D MRI scan was re-acquired for qualitative visualization of the volumetric infusate delivery region. Semi-automated 3D segmentation of Gd signal was later performed to visualize infusate placement in each individual subject; for further details, refer to Supplemental Materials and Supplemental Figure 2.

After the infusions were complete the animal was transported back to the surgical suite. The trajectory guides were removed, and the incision was closed in layers before the animal was allowed to recover from anesthesia. Animals were given buprenorphine twice on the day following the surgery (0.01-0.03 mg/kg, IM). Cefazolin (20-25 mg/kg, IM or IV) or Cephalexin (20-25 mg/kg, oral (PO)) was given twice daily for five days after surgery to prevent infection.

#### Viral vector

Five monkeys received bilateral infusions of the DREADD viral vector AAV5-hSyn-HA-hM3Dq obtained from Addgene (Watertown, MA; lot # v46636; titer 1 x 10^13^ genome copies/ml) that was prepared with construct pOTTC1596 (pAAV SYN1 HA-hM3D(Gq)).

### [^11^C]DCZ Radiosynthesis

The [^11^C]DCZ was synthesized similar to published methods (Nagai et al., 2020) with modifications. The radiolabeling synthon, [^11^C]methyl triflate was produced using the gas phase method, starting with the production of [^11^C]methane via the ^14^N(*p,*α)^11^C nuclear reaction using a 90/10 H_2_/N_2_ target (Jewett, 1992). The [^11^C]methane was converted to [^11^C]methyl iodide through gas phase iodination with iodine vapor in a heated quartz tube using a recirculation process (Larsen et al., 1997). [^11^C]MeI was then passed through a heated glass tube containing silver triflate for the production of [^11^C]methyl triflate (Jewett, 1992). The [^11^C]MeOTf was slowly bubbled into a reaction vial containing 0.2mg 11-(1-Piperazinyl)-5*H*-dibenzo[*b*,*e*][1,4]diazepine precursor (Tocris, Bio-Techne Corp., Minneapolis, MN) and 0.3mL anhydrous MeCN (ThermoFisher Scientific, Waltham, MA). The radiolabeling reaction occurred at room temperature for 5 minutes, after which 0.5 mL mobile phase was added to the vial and the entire solution injected onto the semi-preparative HPLC system. The semi-preparative HPLC system consists of a Waters 515 HPLC pump (5 mL/min), Waters 2489 UV Detector (254 nm), Phenomenex Gemini NX C-18 column (250 mm x 10 mm), and a mobile phase composed of MeCN/H_2_O/Et_3_N (40/60/0.1%) (Decon Labs, ThermoFisher Scientific, Waltham, MA). The peak corresponding to [^11^C]DCZ was collected and diluted in 60 mL water and trapped on a tC18 light cartridge (Waters, Milford, MA) which was then rinsed with 10 mL 0.002N HCl (Fisher). The [^11^C]DCZ compound was then eluted from the cartridge using 0.5 mL EtOH and 10 mL 0.9% sodium chloride, USP (Hospira, Lake Forest, IL) through a Millex-FG membrane filter (MilliporeSigma, Darmstadt, Germany). The purity of the tracer was verified with DCZ standard using a Phenomenex Luna C18 column (200 mm x 4.6 mm) and mobile phase of MeCN/H_2_O/Et_3_N (40/60/0.1%) with a flow rate of 2 mL/min. No measurable radiochemical or chemical impurities were observed. The specific activity was 50.1 ±20.1 mCi/nmol at end of synthesis (N = 5.)

### PET Scanning

[^11^C]DCZ PET scanning was performed in 5 subjects (3 hM3Dq subjects and 2 unoperated controls), after, on average, 700 (± 136) days post-surgery of the hM3Dq subjects. The subjects were anesthetized with ketamine (15 mg/kg, IM), given atropine sulfate (0.04 mg/kg, IM) to depress salivary secretion, intubated and received isoflurane anesthesia (1–1.5%, IT), and placed in the Focus 220 microPET scanner in the prone position with the head facing downward and secured in a custom-made head holder. Throughout the experimental procedure vital signs (heart rate, respiration, oxygen saturation, end tidal CO_2_, and body temperature) were continuously monitored. To maintain fluids and electrolytes, Plasmalyte (up to 10 mg/kg/hr, IV) was administered. A transmission scan was acquired for 518 seconds using a ^57^Co rotating rod source to correct for the scatter and attenuation of radiation in the tissue. The PET scan was initiated with the injection of the [^11^C]DCZ (N=5; 4.9mCi ± 0.18 mCi; 0.07 ± 0.04 micrograms DCZ). The list mode emission data was acquired for a total of 90 minutes and binned into a time series of 22 frames: 2 x 1 min, 4 x 2 min, 16 x 5 min. The PET data were reconstructed using filtered back-projection and a ramp filter into a matrix size of 128 x 128 x 95 with voxel dimensions of 0.959mm x 0.959mm x 0.796mm.

### [^11^C]DCZ Binding Image Analysis

To quantify specific binding of [^11^C]DCZ, a parameter representing radioligand-receptor binding was calculated to serve as an index of neuroreceptor density. The distribution volume ratio (DVR) was estimated from the PET time series for each voxel in this parametric image, and DVR images were calculated using the SRTM2 algorithm (Gunn et al., 1997; Wu & Carson, 2002), with the cerebellar grey matter as the reference region of negligible specific ligand-receptor binding. The DVR images were spatially transformed into a common T1-weighted MRI template space made from 592 rhesus monkeys (Fox et al., 2015a) and blurred with a 2mm FWHM smoothing kernel using Advanced Normalization Tools (ANTS; Avants et al., 2010, 2011).

For the group analysis, to control for individual variation in baseline [^11^C]DCZ binding observed between subjects, the DVR images were scaled such that specific binding signal in the medial prefrontal cortex was matched across all subjects, following previously published methods (Roseboom et al., 2021). This was done to normalize for overall global differences in DCZ-specific receptor expression across subjects, and the medial prefrontal cortex was selected because this region revealed the highest and most uniform values of [^11^C]DCZ DVR across subjects. The image from the average of the two control subjects was then subtracted from the average of the three hM3Dq DREADD subjects using FSL (FMRIB).

In a separate analysis to understand the anterior-posterior extent of [^11^C]DCZ binding, the DVR images from the three hM3Dq subjects were aligned to template space and to each other using only linear (affine) transformations to preserve the spatial relations among the injection sites. In the first step, subject E1’s [^11^C]DCZ image was aligned to the MRI template with FLIRT (FMRIB’s Linear Image Registration Tool) using a between-modality cost function (Normalized Mutual Information). Next, subject E3’s [^11^C]DCZ image was aligned to subject E1’s [^11^C]DCZ image in template space using a within-modality cost function (Normalized Correlation). The same was done for subject E5’s [^11^C]DCZ image. All three aligned images were then blurred with a 1mm FWHM smoothing kernel using ANTS.

### Immunohistochemical procedures

Time between surgery and necropsy was 667 days for subject E1, 701 days for subject E5, and 807 days for subject E3. Animals were euthanized by transcardial perfusion with heparinized PBS at room temperature followed by fixation with 4% paraformaldehyde (PFA) in PBS at 4°C. In the case of subject E3, which was also used for electron microscopy studies, perfusion was performed using ice-cold, oxygenated Ringer’s buffer followed by 4% PFA containing 0.1% glutaraldehyde in 0.2 M phosphate buffer, pH 7.4. Brains were removed and fixation was continued in the perfusion solution overnight at 4 °C. The brains were placed in a brain block and cut into slabs (approximately 14 mm thick) that were subsequently processed through increasing concentrations of sucrose: 10%, 20%, and 30%. Slabs containing the amygdala, thalamus, or frontal cortex were frozen and cut at 40 microns on a cryostat (CryoStar NX50, ThermoFisher Scientific). Tissue sections were stored in cryoprotectant (40 mM potassium phosphate/11mM sodium phosphate buffer, pH 7.2, containing 30% ethylene glycol and 30% sucrose) at -20°C until use.

Chromogenic labeling of HA-tag was performed to observe hM3Dq-HA expression within the amygdala, as well as in regions with which the amygdala shares monosynaptic connections (i.e., orbitofrontal cortex, bed nucleus of the stria terminalis, thalamus, etc.) Tissue sections were washed with PBS at room temperature to remove cryoprotectant, and all subsequent incubations were carried out at room temperature in PBS containing 0.3% Triton X-100. Three 5-min washes with PBS were performed in between each incubation step. To block endogenous peroxidase activity, tissue sections were incubated in PBS containing 6% hydrogen peroxide for 30 minutes. Sections were then blocked in 5% normal goat serum (Cat # S-1000; Vector Laboratories, Burlingame, CA, USA) for 1 hour. This was followed by an overnight incubation with a 1:400 dilution of rabbit monoclonal HA-tag antibody, and then a 1-hour incubation with a 1:250 dilution of peroxidase-conjugated goat anti-rabbit immunoglobulin G (IgG) secondary antibody. See Supplemental Table 2 for details on antibodies utilized. A 2-minute incubation with ImmPACT DAB peroxidase substrate (Cat # SK-4105; Vector Laboratories) was then used for chromogenic visualization. Sections were mounted on Superfrost Plus microscope slides, which were air-dried, dehydrated in 95% and then 100% ethanol, and placed in xylene for 5 min before coverslipping with DPX mounting medium. Images were captured using a DM6000B light microscope (Leica Microsystems, Buffalo Grove, IL, USA).

Immunofluorescent co-labelling of HA-tag, neuronal nuclei (NeuN), and/or glial fibrillary acidic protein (GFAP) was performed to characterize expression of hM3Dq-HA in relation to neurons and astroglia. Tissue sections were washed with PBS at room temperature to remove cryoprotectant, and all subsequent incubations were carried out at room temperature in PBS containing 0.3% Triton X-100; three 5-minute washes with PBS were performed in between each incubation step. Prior to incubation with a primary antibody, tissue sections were first blocked in either 5% normal goat serum (Cat # S-1000; Vector Laboratories, Burlingame, CA) or 5% normal donkey serum (Cat # 017-000-121; Jackson ImmunoResearch, Laboratories, Inc., West Grove, PA) for one hour. This was followed by a 24-hour incubation with the first primary antibody and then a one-hour incubation with an Alexa Fluor secondary antibody. Subsequent incubations in blocking buffer, primary antibody solution, and secondary antibody solution were carried out for each additional marker of interest, such that the following combinations were performed: HA-tag and NeuN, GFAP and NeuN, GFAP and HA-tag and NeuN. Cellular nuclei were stained using a 1:10000 dilution of 4’,6-diamidino-2-phenylindole (DAPI) for 5 minutes. Sections were mounted on Superfrost Plus microscope slides using ProLong Gold Antifade Mountant (Cat # P36930; ThermoFisher Scientific).

### Imaging and Stereological Quantification of Expression in the Central Nucleus

For each subject examined, 5 coronal sections spanning the anterior-posterior plane of the Ce in one hemisphere (left or right was randomly chosen) were utilized for analysis of HA-tag and NeuN co-expression. Sections immediately adjacent to these were processed for acetylcholinesterase (AChE), a cholinergic marker that was used to identify nuclei within the amygdala (Amaral and Bassett, 1989; Paxinos et al., 2009). Fluorescent image stacks were acquired using a Nikon A1R-Si+ confocal microscope with a pan fluor 40x oil objective (1.30 N.A.) and Nikon NIS-Elements software. All sections were imaged using the same acquisition settings, including magnification, laser power, camera gain, offset, pinhole size, and scan speed. To evaluate neuronal expression of hM3Dq-HA, standard unbiased stereological counting of HA and NeuN immunopositive cells was accomplished with the optical fractionator workflow in Stereo Investigator software (MBF Bioscience). Counting parameters were similar to those utilized in stereological assessment of hM4Di-HA expression in the Ce, described in (Roseboom et al. 2021); for further details concerning the stereological parameters used, refer to Supplemental Table 3. Cells that were immunoreactive for NeuN and HA-tag were identified using the soma as the counting target. Cells with a well-defined soma and nucleus were counted if they were within or intersecting with the dissector frame and not intersecting with lines of exclusion. Counting was performed by a single trained investigator.

### Electron microscopy

A slab of tissue from subject E3 containing the amygdala was processed for electron microscopy. The slab was cut in a vibratome at 50 µm thickness and sections were stored in anti-freeze solution until ready to use. The sections selected for EM immunoperoxidase were treated with 1% sodium borohydride (20 min), placed in a cryoprotectant solution (PB, 0.05 M, pH 7.4, containing 25% sucrose /10% glycerol) and frozen at -80° C (each of these steps lasted 20 min).

Sections were thawed and washed in PBS. Then, to block non-specific sites, the sections were immersed in PBS containing 5% non-fat dry milk (20 min). Sections were then incubated in antibodies against HA-tag (same as used in light and fluorescent microscopy studies described above, see Supplemental Table 2), for 48 hours. This was followed by incubation in biotinylated goat anti-rabbit secondary antibodies (1:200, Vector Laboratories (Burlingame, CA), catalog number BA-1000, RRID AB-2313606) for 90 min, and in avidin-biotin-peroxidase complex (ABC) solution (1:200; Vectastain standard kit, Vector) for 90 min. The staining was revealed by exposure to 0.025% 3-3’-diaminobenzidine tetrahydrochloride (DAB, Sigma-Aldrich, St. Louis, MO), 0.01M Imidazole (Fisher Scientific, Pittsburgh, PA) and 0.006% H2O2 for 10 min. Next, the sections were rinsed in PB (0.1 M, pH 7.4) and treated with 0.5% osmium tetroxide (OsO4) for 10 min and returned to PB. Sections were then exposed to increasing concentrations of ethanol. To increase EM contrast, 1% uranyl acetate was added to the 70% ethanol solution (10 min in the dark). Sections were placed in propylene oxide, followed by tissue embedding with an epoxy resin (Durcupan; Fluka, Buchs, Switzerland) for 12 hr. The resin-embedded sections were cured at 60 LJC (48 hours). All incubations were done at room temperature. Control sections were incubated without the primary antibodies.

Sections were examined under the light microscope, and blocks of tissue in the regions of interest were cut and glued onto resin blocks. Blocks containing regions of interest were obtained. Ultrathin sections (60 nm) were obtained from these blocks using an ultramicrotome (Ultracut model T2, Leica, Nussloch, Germany) and mounted onto pioloform-coated copper grids. The sections were exposed for 5 minutes to lead citrate to increase their contrast for EM visualization. Blocks were obtained in the areas of interest, and then cut into ultrathin 60 nm sections (Ultracut T2, Leica).

In addition to the immunoperoxidase EM staining, some sections were treated with immunogold staining, because the immunogold method provides a much higher spatial resolution to identify the localization of the antigen of interest (Galvan et al., 2006). Sections selected for immunogold were treated similar as above, with the following changes. To block non-specific sites, the sections were immersed in PBS containing 5% non-fat dry milk (20 min) and washed three times in Tris-buffered saline (TBS)-gelatin (Tris 0.02 M, NaCl 0.15 M and 0.1% cold water fish gelatin, 10 min each wash). Sections were incubated with the primary antibodies against HA-Tag for 24 hours. Then, sections were rinsed in TBS-gelatin and incubation in gold-conjugated secondary antibodies for 2 hours. Primary and secondary antibodies were diluted in TBS-gelatin buffer containing 1% dry milk. The sections were then washed with TBS-gelatin and 2% acetate buffer (pH 7.0) and the gold labeling was intensified with silver (HQ silver; Nanoprobes) for 9-13 min in the dark. The silver intensification was terminated with acetate buffer rinses. The tissue was then post-fixated with osmium tetroxide and embedded in resin. Blocks were cut in the regions of interest and ultrathin sections were obtained as described above.

Sections were examined with an electron microscope (JEOL model 1011, Peabody, MA) at 40,000-60,000X, and electron micrographs were obtained in areas containing immunolabeled elements using a digital camera (DualView 300W; Gatan, Inc., Pleasanton, CA) controlled by DigitalMicrograph software (Gatan, version 3.11.1). Micrographs were examined using ImageJ (Rasband, 1997; Schneider et al., 2012).

### Analysis of material

Electron micrographs were collected from areas with optimal ultrastructural preservation of brain tissue. The micrographs contained at least 112 immunoperoxidase-labeled elements from each block (total area examined ranged from 509 to 904 micrometers^2^). Two blocks of tissue each were analyzed for the lateral division (CeL) and the medial division (CeM) of the Ce, and one block of tissue each was analyzed for the dorsal basal and accessory basal nuclei. The subcellular compartments containing immunoperoxidase were categorized into dendrites, axonal terminals, unmyelinated axons or astrocytic processes based on ultrastructural characteristics as described by Peters et al. (Peters et al., 1991).

Elements were considered to be HA-tag positive if they contained dense amorphous deposits characteristic of immunoperoxidase. The relative proportion of each type of immunolabeled element was calculated for each nucleus; for the CeL and CeM, these proportions were averaged across the two blocks of tissue examined. See Supplemental Table 5 for information regarding the number and distribution of elements quantified in this analysis.

## Results

### *In vivo* imaging to verify infusion target and DREADD expression

To verify the location of the infusion during surgery, a gadolinium contrast agent was added to the viral infusate. iMRI voxels from post-infusion scans showing increased gadolinium signal were segmented, and this segmentation was overlaid on the pre-infusion scan acquired at the start of the iMRI procedure, shown in Figure 2. Infusate was observed in the bilateral amygdala of all subjects, with some variation in its anterior-posterior extent. To demonstrate the localization of hM3Dq *in vivo*, [^11^C]DCZ PET imaging was performed in three of the hM3Dq subjects and two of the control subjects approximately 681 (± 107) days post-surgery. Scaled DVR images were averaged for each group, and the mean image from the control group was subtracted from the mean image of the hM3Dq group. Figure 3 depicts the image of the difference between the hM3Dq-HA and Control groups, thresholded at ≥40% and overlaid on a rhesus monkey brain MRI template. The resulting elevated [^11^C]DCZ signal was detected exclusively in the bilateral amygdala region. The voxel with the average peak difference was observed in the right amygdala, reaching a 52% increase in binding.

**Figure 2.**
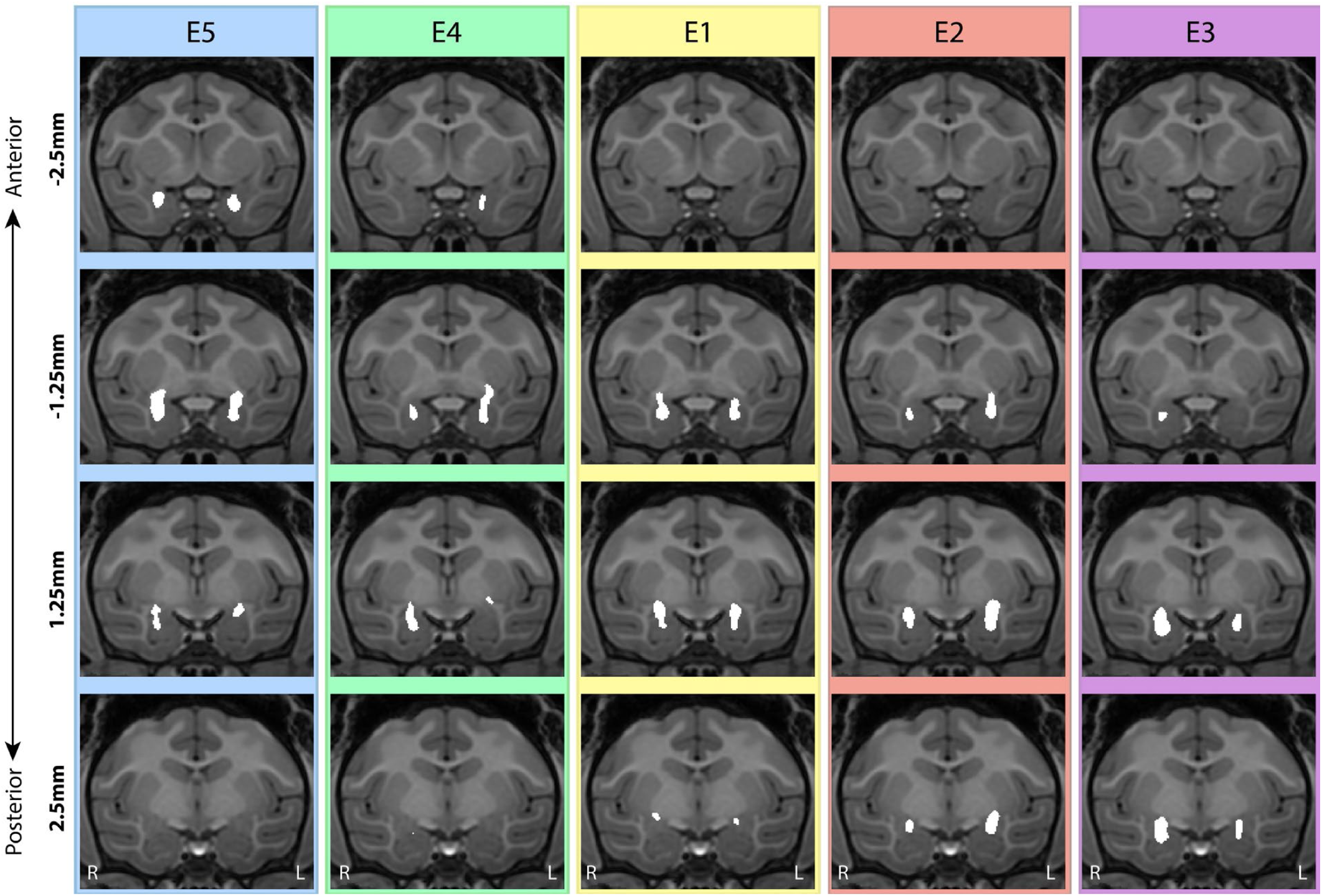
*In vivo* MRI visualization of AAV5-hSyn-HA-hM3Dq infusion. Voxels indicative of gadolinium signal were segmented (shown in white) and overlaid on a pre-infusion template for each subject. These voxels were observed in the bilateral amygdala of each subject, with some variation in anterior-posterior placement between subjects, which are arranged in order of this variation, e.g., subject E5 had the most anterior infusion, and subject E3 had the most posterior infusion. The anterior-posterior measurements are relative to the posterior edge of the anterior commissure (0.0 mm). R, right hemisphere; L, left hemisphere.

**Figure 3.**
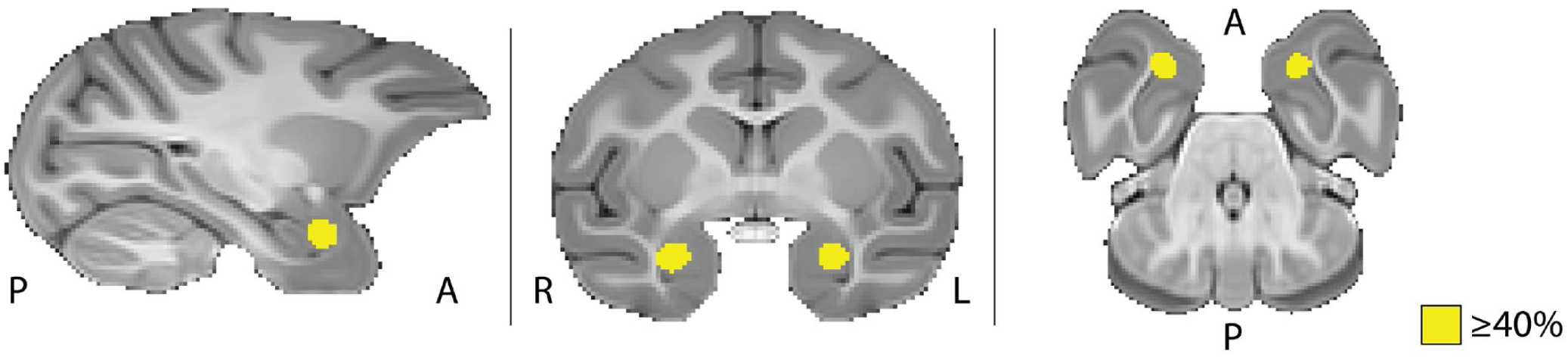
[^11^C]DCZ binding in the amygdala demonstrates hM3Dq-HA expression *in vivo*. The average difference (hM3Dq-HA minus control; ≥40%), reveals greater [^11^C]DCZ binding bilaterally in the amygdala of hM3Dq-HA subjects relative to controls. Sagittal (left), coronal (middle), and axial (right) views are shown. A, anterior; P, posterior; R, right hemisphere; L, left hemisphere.

### Histological assessment of hM3Dq-HA expression

#### Regional localization of hM3Dq-HA

The three hM3Dq subjects discussed above were also used to verify hM3Dq-HA expression histologically. Figure 4A shows the viral infusion, [^11^C]DCZ binding (DVR), and HA-DAB images (at 1.25X magnification) for subject E5 (Supplemental Figure 3 shows these same measures for subjects E1 and E3). It is notable that, even with the 681 (± 107) day interval between the acquisition of the gadolinium images and the [^11^C]DCZ PET imaging, there is substantial correspondence within animals in relation to the variation in anterior-posterior extent of these signals. Furthermore, intensity of [^11^C]DCZ binding in the amygdala was consistent with the relative amount of HA-tag staining observed, such that the regions of high binding intensity were also highly HA-immunoreactive within each subject.

**Figure 4.**
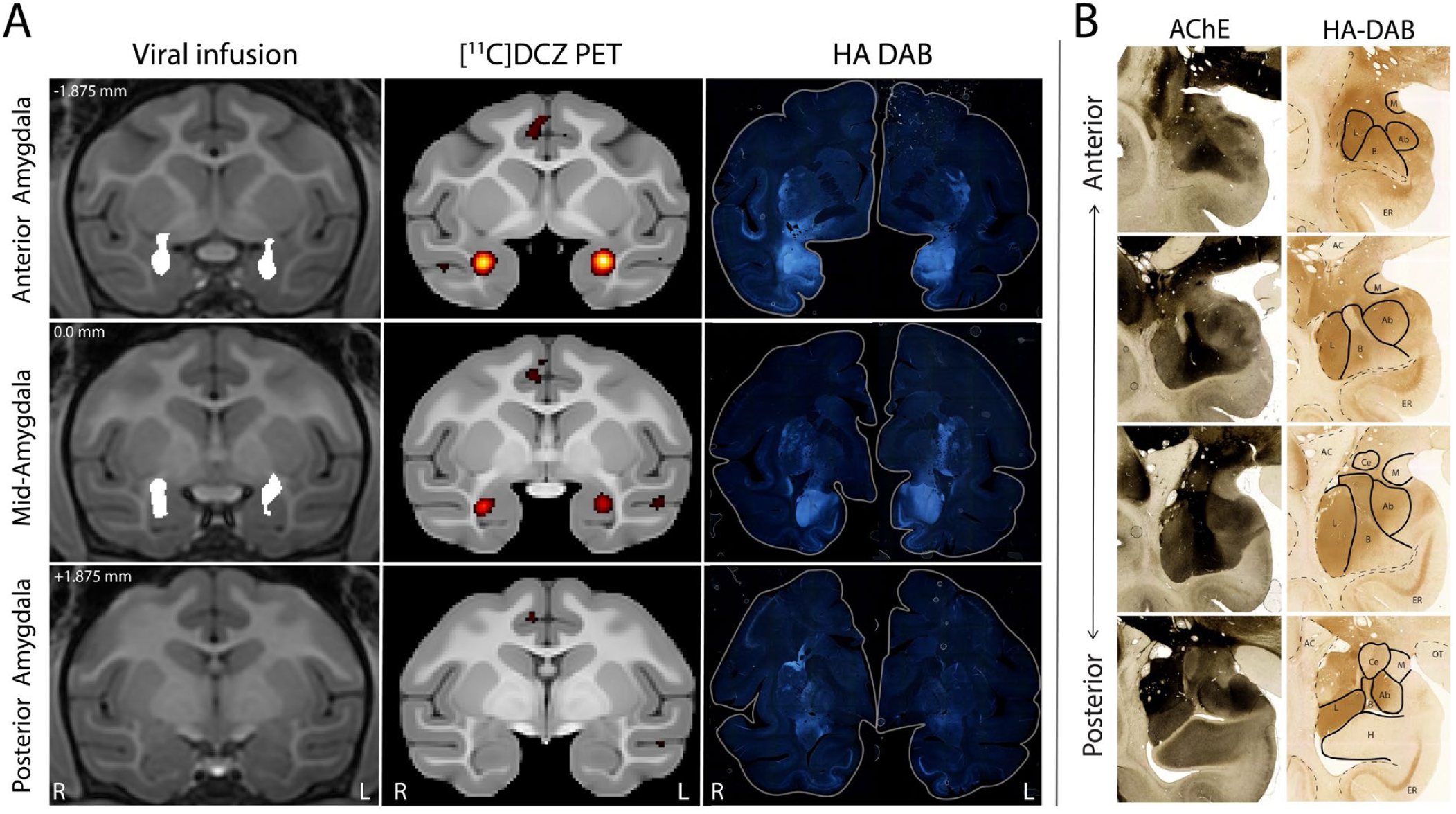
Detection of hM3Dq-HA expression in the NHP amygdala across multimodal measures. A) Images showing viral infusion, elevated [^11^C]DCZ PET binding, and immunohistochemical detection of HA-tag are juxtaposed to demonstrate correspondence of these measures within subject E5. Viral infusion images show segmented voxels (white) indicative of gadolinium signal, which was used to monitor infusion of AAV5-hSyn-HA-hM3Dq. [^11^C]DCZ PET imaging (thresholded at ≥80% of subject’s peak binding value) demonstrates binding in the amygdala. Immunoperoxidase DAB labeling was used to visualize HA-tag post-mortem, shown in full coronal slice view. Zero (0.0 mm) corresponds to the posterior edge of the anterior commissure. R, right hemisphere; L, left hemisphere. **B)** Tissue sections from the right hemisphere of subject E5 were stained using acetylcholinesterase for anatomical identification of amygdala nuclei, which are outlined in solid black on the HA-DAB images. In adjacent sections, immunoperoxidase DAB labeling was used to visualize HA-tag. HA-tag immunoreactivity can also be seen in the entorhinal cortex. Ab, Accessory basal nucleus; AC, anterior commissure; B, basal nucleus; Ce, central nucleus; ER, entorhinal cortex; H, hippocampus; L, lateral nucleus; M, medial nuclei; OT, optic tract.

Within these subjects, HA+ cell bodies and neuropil were found throughout all nuclei of the amygdala. The greatest density of HA-immunoreactivity was present in the basolateral nuclei (BLA, including the accessory basal, basal, and lateral nuclei), where HA+ neuropil was extremely prominent, while HA+ neuropil was considerably less prominent in the Ce (an example of this in subject E3 can be seen in Figure 5A). Little to no HA immunoreactivity was observed in the medial nucleus. Figure 4B displays the HA+ immunoreactivity throughout the amygdala of one animal (subject E5), which is representative of the pattern described above.

**Figure 5.**
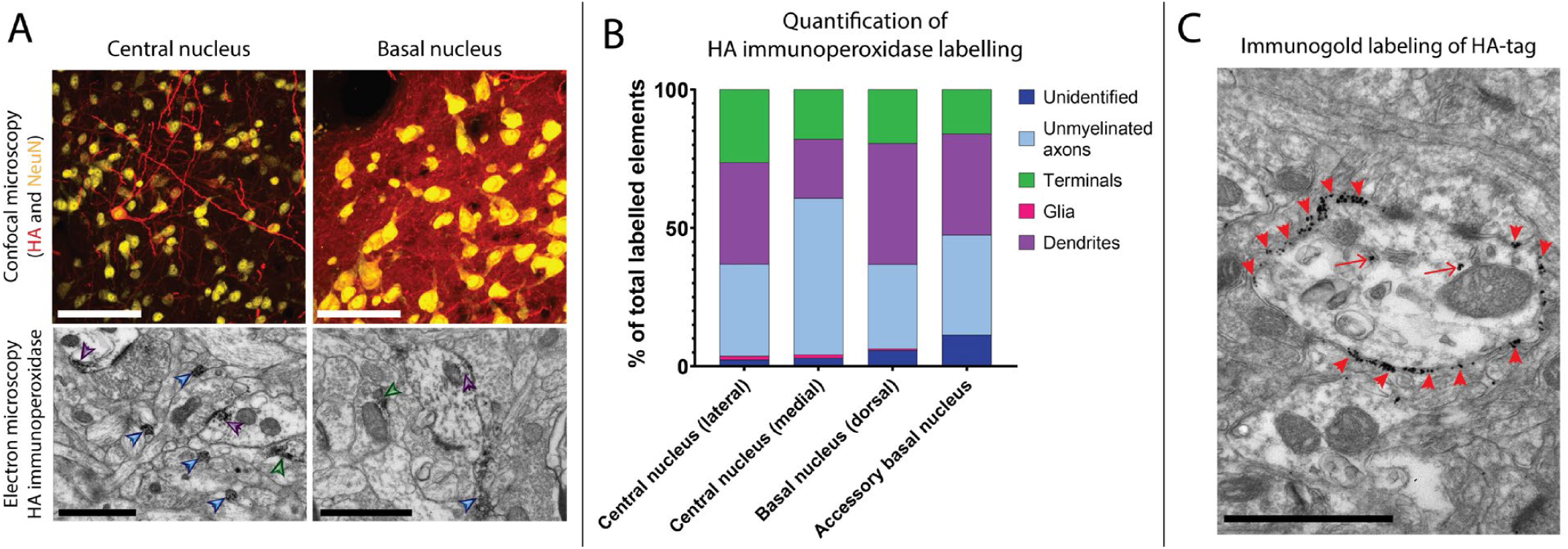
Cellular and ultrastructural localization of hM3Dq-HA in the NHP amygdala. In the right hemisphere of subject E3, ***A*)** fluorescent staining of HA tag (red) is observed in neuronal cell bodies (stained with NeuN in yellow) as well as neuropil, with HA-tag expression generally appearing more abundant in the basolateral nuclei (including the basal nucleus) than in the central nucleus. At the ultrastructural level, HA-immunoperoxidase labeling is found in both pre- and post-synaptic neuronal elements, with examples shown in dendrites (purple arrowheads), terminals (green arrowheads), and unmyelinated axons (light blue arrowheads). Immunoperoxidase labeling was then quantified across these elements **(B)**. Examples of immunogold particles indicating hM3Dq-HA are shown in the BLA **(C)**, where red arrowheads point toward examples of immunogold particles on the plasma membrane, and red arrows point toward immunogold particles in the intracellular space. Scalebars are 100 μm (white, confocal microscopy) or 1.0 μm (black, electron microscopy in panels A and C).

Although numerous cell bodies were immunoreactive for HA-tag in the BLA, these could not be accurately quantitated because of the density of HA-tag expression in the surrounding and overlying neuropil. In comparison, because there was much lower expression in the Ce, immunopositive cell bodies were able to be quantitated using stereological estimates, revealing that approximately 6% (on average) of Ce neurons expressed hM3Dq-HA (transduction efficiency range 4 – 9%; see Supplemental Table 4 for details). The highest percentage of Ce neuronal expression (9%) was observed in subject E3. This may in part be due to superior preservation of cellular morphology in this subject, as this tissue was fixed with the addition of glutaraldehyde for examination using electron microscopy (see “Electron Microscopy” results below). Additionally, some tissue damage in the Ce of subject E1 was also apparent, as indicated by an absence of NeuN and HA-tag immunoreactivity in an anterior portion of the Ce. This portion was excluded from the stereological sampling conducted within this subject.

Some immunohistochemical labeling of HA-tag was observed outside of the amygdala, including regions that are known to share monosynaptic connections with the amygdala. In all three subjects examined, HA+ cell bodies and neuropil were observed in the entorhinal and insular cortices (visible in Figure 4), as well as in the hippocampus and putamen. Sparse labeling of HA-tag was also found in the pregenual ACC (Area 25) and orbitofrontal cortex (OFC) of subject E3. This extra-amygdalar expression was not detected by [^11^C]DCZ PET.

Additionally, in all of the subjects examined, a small area that was devoid of NeuN and HA immunoreactivity was found in the region of the injection site—in particular, the anterior portion of the dorsal basal nucleus. Immunohistochemical characterization of this region revealed a high presence of glial fibrillary acidic protein (GFAP), indicative of activated astroglia (see Supplemental Figure 4). Furthermore, no colocalization of HA tag and GFAP was observed, confirming that hM3Dq-HA expression was not occurring in astroglia (also shown in Supplemental Figure 4).

#### Cellular and subcellular localization of hM3Dq-HA

Double labelling with the neuronal marker NeuN confirmed that hM3Dq was selectively expressed in neurons (see Figure 5A, top panel). At the ultrastructural level, peroxidase immunoreactivity for HA-tag was observed in unmyelinated axons, axon terminals, and dendrites throughout the amygdala, indicating that expression of hM3Dq-HA occurs in both pre- and post-synaptic neuronal elements (Figure 5A, bottom panel, and 5B). Additionally, immunogold labeling (Figure 5C) was used to examine the localization of hM3Dq-HA in relation to plasma membranes. The large majority of gold particles were found to be membrane-bound, indicating that hM3Dq-HA is successfully transported and incorporated into the plasma membrane of NHP amygdala neurons.

### Experiment 1: Clozapine-induced hM3Dq-HA activation enhances anxiety-related responses

#### Anxiety-related responses during the Alone and NEC conditions

Based on previous work (Kalin & Shelton, 2003; Roseboom et al., 2021), we hypothesized that DREADD-mediated amygdala activation would result in enhanced anxiety-related responding across contexts after clozapine treatment. Confirming this hypothesis, the results revealed a significant 3-way interaction of Group (control versus hM3Dq) x Treatment (vehicle versus clozapine) x Pre/Post (pre-surgical versus post-surgical testing) for freezing duration (F_1,376_ = 4.218, p < 0.05) and locomotion (F_1,376_ = 15.247, p < 0.001), while controlling for plasma clozapine concentrations. These 3-way interactions demonstrate cross-context effects of DREADD activation, which are consistent with the lack of significant 4-way interactions of Group x Treatment x Pre/Post x Condition (Alone versus NEC) for freezing and locomotion. No significant 3- or 4-way interactions were observed for coo-vocalizations. *Post-hoc* testing of the significant 3-way freezing interaction revealed during the postsurgical period that clozapine treatment, relative to vehicle, resulted in significant increases in freezing in the hM3Dq subjects (t(154) = 4.083, p < 0.001; see Figure 6). In contrast, significant decreases in freezing in the control subjects were observed in response to clozapine treatment during the postsurgical period (t(154) = -3.573, p < 0.001). For locomotion, *post-hoc* testing revealed significant clozapine-induced decreases in the hM3Dq subjects (t(154) = -7.676, p < 0.001) and significant increases in the control subjects (t(154) = 3.226, p < 0.001, see Supplemental Figure 7). Refer to Supplemental Table 6 for full statistical results. See Supplemental Figures 5 and 6 for untransformed freezing and locomotion data.

**Figure 6.**
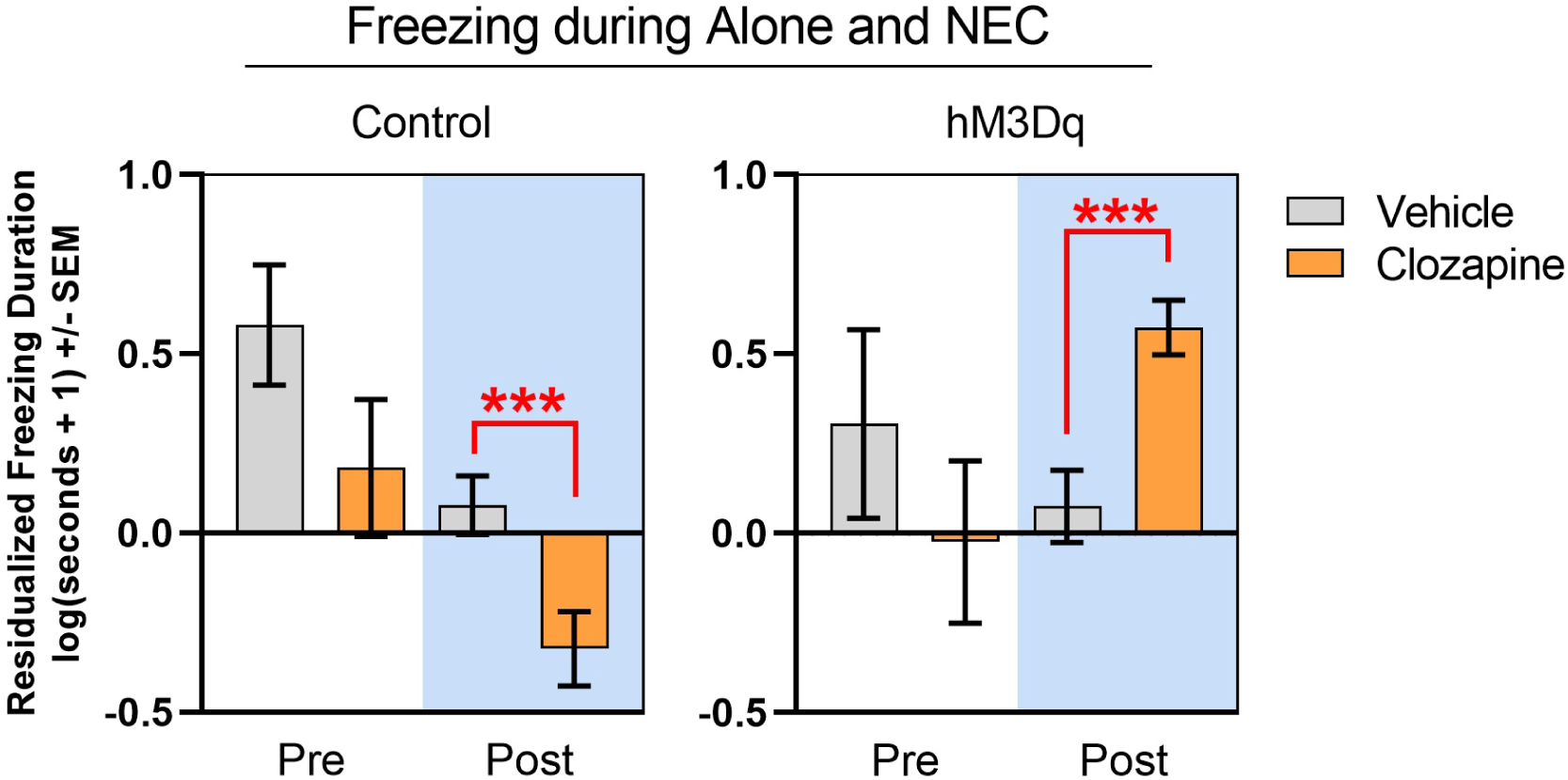
**hM3Dq-mediated amygdala activation increases freezing behavior during the Alone and NEC conditions**. Graphs show mean log-transformed freezing durations (residualized for plasma clozapine concentrations) across the Alone and NEC conditions of the HIP. Data following vehicle or clozapine administration are shown for both the control and hM3Dq groups during pre-surgical (white background) and post-surgical (blue background) testing. A significant Group x Treatment x Pre/Post interaction was observed (F_1,376_ = 4.218, p < 0.05). ***p<0.001

#### Anxiety-related responses during the Stare condition

Data from the Stare condition was collected only after surgery. Experimenter hostility and bark vocalizations are the prominent responses observed during the Stare condition, however, no significant Group x Treatment interactions were observed for either of these behaviors. The analysis of freezing behavior, which typically does not occur during Stare, revealed findings that are consistent with the above analyses of freezing occurring in the Alone and NEC conditions. hM3Dq activation resulted in a significant 2-way interaction for freezing (F_1,68_ = 11.271, p = 0.001), and *post-hoc* testing revealed that hM3Dq subjects froze more (t(34) = 4.941, p < 0.001) following clozapine treatment, relative to vehicle, while no significant changes were observed in Control subjects (see Figure 7). Analysis of locomotion data also revealed a significant 2-way interaction (F_1,68_ = 11.762, p = 0.001, see Supplemental Figure 8). *Post hoc* testing revealed that hM3Dq subjects locomoted less (t(34) = -3.089, p < 0.01) following clozapine treatment, relative to vehicle, while no significant changes were observed in Control subjects. No significant interactions were observed for coo-vocalizations. Refer to Supplemental Table 7 for full statistical results.

**Figure 7.**
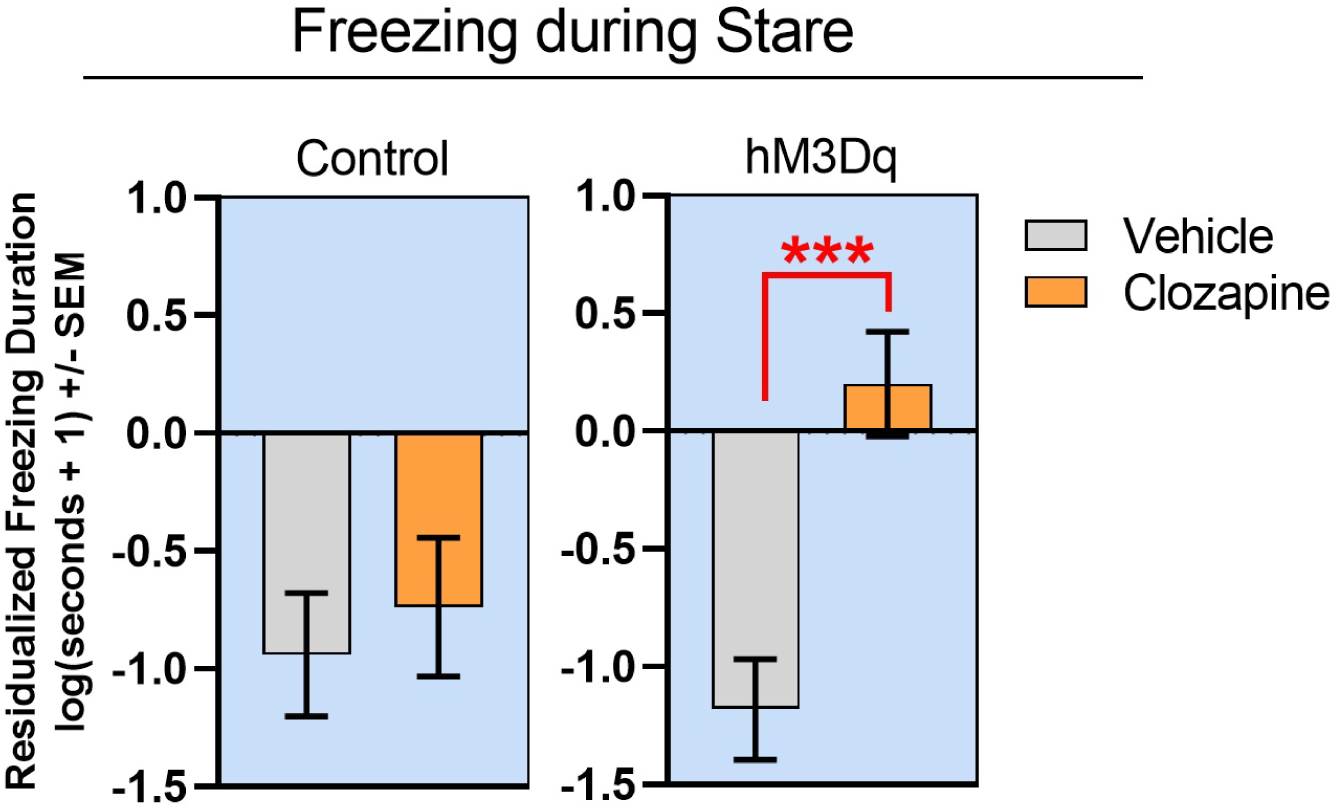
**hM3Dq-mediated amygdala activation increases freezing behavior during the Stare condition**. Mean log-transformed freezing durations (residualized for plasma clozapine concentrations) following vehicle or clozapine administration are shown for both the control and hM3Dq groups during the Stare condition. A significant Group x Treatment interaction was observed (F_1,68_ = 11.271, p = 0.001). (***p<0.001)

### Experiment 2: Anxiety-related responses continue to be affected by DREADD-mediated amygdala activation two years following viral vector transduction

To replicate the effects observed in Experiment 1 and to assess the extent to which DREADDs remain responsive over an extended period of time, behavioral testing of the animals was repeated separately with clozapine, DCZ, and vehicle in the full HIP (Alone, NEC, and Stare) approximately two years after surgery. We used DCZ in addition to clozapine as DCZ is an effective DREADD-actuator with greater selectivity for DREADDs than clozapine (Fujimoto et al., 2021; Nagai et al., 2020; Upright & Baxter, 2020; Yan et al., 2021).

Results following clozapine or DCZ treatment were generally consistent with the effects observed in Experiment 1. Relative to vehicle, increased freezing was observed in hM3Dq subjects following clozapine administration, such that there was a significant Group x Treatment interaction (F_1,142_ = 13.366, p < 0.001; see Figure 8), again suggesting that the anxiogenic effect persisted across contexts. Additionally, a significant 2-way interaction for freezing was found following DCZ administration (F_1,142_ = 4.756, p < 0.05; see Figure 9). Decreased locomotion occurred in hM3Dq subjects following clozapine administration across all contexts, as characterized by a significant Group x Treatment interaction (F_1,142_ = 16.814, p < 0.001; see Supplemental Figure 11). In relation to DCZ administration, a significant 3-way interaction (F_2,142_ = 3.428, p < 0.05) was also detected, as characterized by decreases in locomotion only during the Alone condition (t(43) = -3.633, p < 0.001; see Supplemental Figure 12). See Supplemental Figures 9 and 10 for untransformed freezing and locomotion data. Unlike the results of Experiment 1, there were some significant effects on coo-vocalizations (Supplemental Figures 13 and 14) and bark vocalizations (Supplemental Figures 15-17) following clozapine and DCZ administration. Refer to Supplemental Tables 8 and 9 for full statistical values.

**Figure 8.**
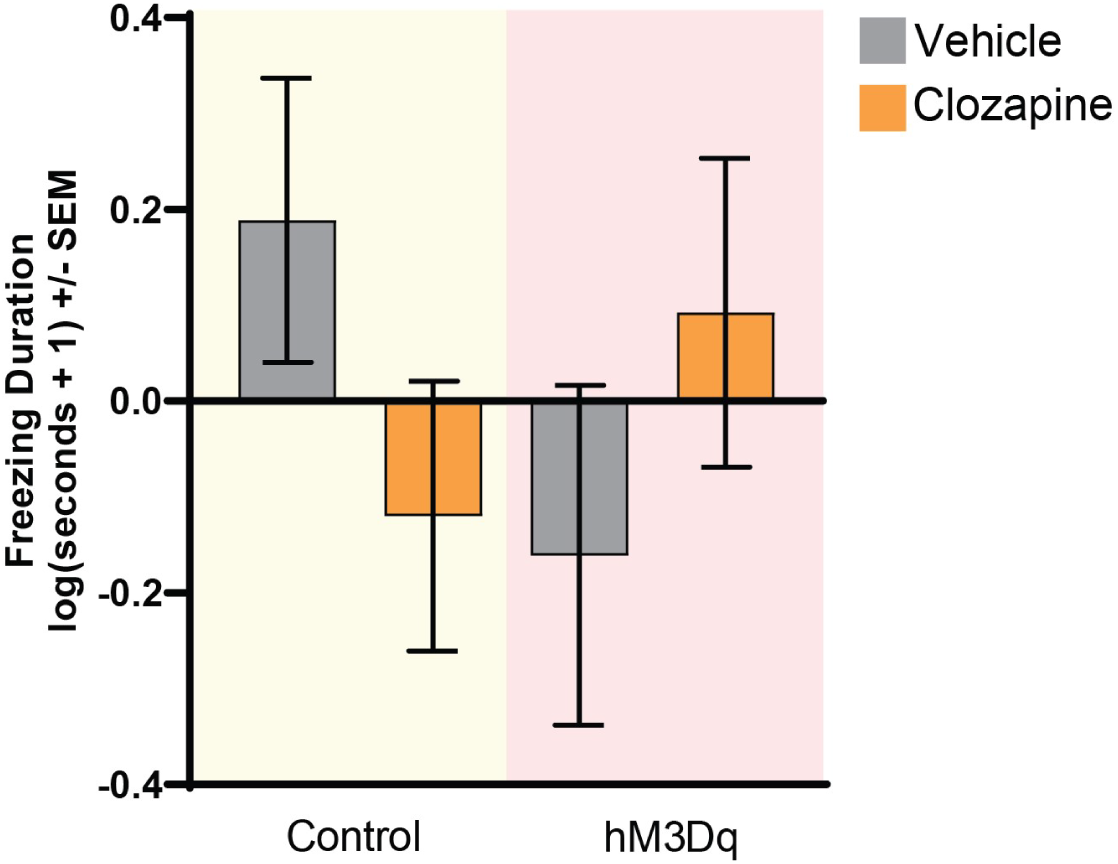
**Clozapine-mediated hM3Dq activation continues to increase anxiety-related freezing after long-term hM3Dq expression**. Mean log-transformed freezing durations following vehicle or clozapine administration are shown for both the control (yellow background) and hM3Dq (red background) groups. The significant Group x Treatment interaction (F_1,142_ = 13.366, p < 0.001) suggests that the effects of DREADD activation occur across contexts. *Post hoc* comparisons revealed no significant treatment-related differences in either group.

**Figure 9.**
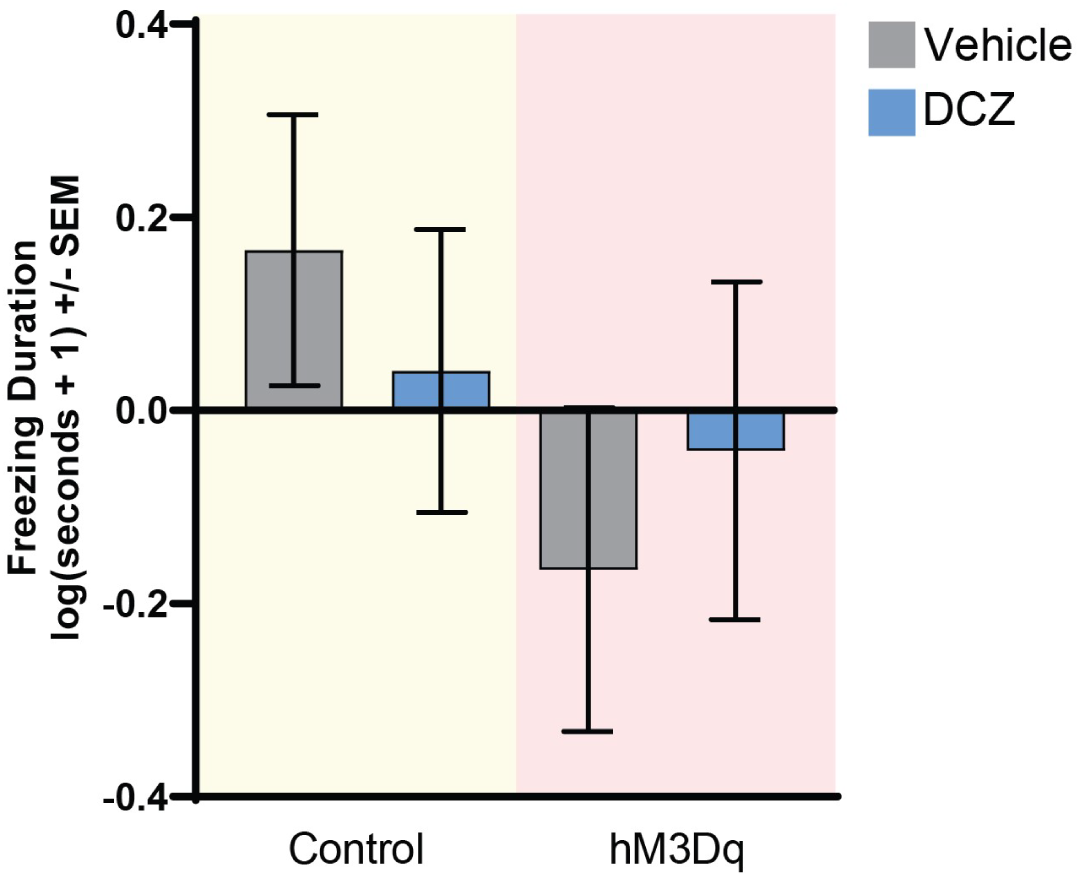
**DCZ-mediated hM3Dq activation increases anxiety-related freezing after long-term hM3Dq expression**. Mean log-transformed freezing durations following vehicle or DCZ administration are shown for both the control (yellow background) and hM3Dq (red background) groups. The significant Group x Treatment interaction (F_1,142_ = 4.756, p < 0.05), suggests that the effects of DREADD activation occur across contexts. *Post hoc* comparisons revealed no significant treatment-related differences in either group.

## Discussion

In this study, we demonstrate that activation of the excitatory hM3Dq DREADD in the NHP amygdala with low-dose clozapine and DCZ is sufficient to increase anxiety-related behaviors, both within and outside of the contexts in which they adaptively occur. We also demonstrate that DREADDs remain functional over the long-term and replicate previous reports demonstrating that [^11^C]DCZ PET imaging can be used to visualize hM3Dq expression *in vivo* (Nagai et al., 2020). [^11^C]DCZ PET imaging allows for the ability to verify DREADD expression in living subjects, which is important for the success of DREADD experiments in NHPs and also constitutes a critical step when considering translation of these methods to humans. Post-mortem immunohistochemical examination determined that hM3Dq-HA was expressed throughout the entire amygdala—most abundantly in neuropil within the basolateral nuclei—and electron microscopy revealed that hM3Dq-HA was predominantly localized to neuronal membranes in both pre- and post-synaptic elements. This complements our previous findings that hM4Di-HA is also predominantly expressed at neuronal membranes (Galvan et al., 2019).

It has been established in numerous studies across species that stimulation of the amygdala can lead to behavioral and autonomic changes (Davis, 1992). In human patients with drug-resistant epilepsy, electrical stimulation of the BLA has been shown to induce physiological alterations in heart rate, blood pressure, and skin conductance (Inman et al., 2020)—responses which are associated with defensive motivation and threat processing (Bradley et al., 2001; Lang & Bradley, 2010). In some cases, emotional states of fear, anxiety, and sadness have also been reported following amygdala electrical stimulation (Lanteaume et al., 2007; Meletti et al., 2006). In macaques, pharmacological activation of the BLA has been shown to lead to reductions in social behavior (Wellman et al., 2016). More recent studies using DREADDs in rodents have demonstrated that activation of hM3Dq-expressing neurons in the BLA is sufficient to induce freezing behavior and impact fear-learning (Sengupta et al., 2016; Yiu et al., 2014; Yoshii et al., 2017). In the present study, we build upon earlier work by demonstrating in a NHP model of pathological anxiety that chemogenetic activation of the amygdala results in increased expression of anxiety-related behaviors. Specifically, by testing the effects of amygdala activation in various threat-related contexts of the HIP, we found that DREADD-induced increases in freezing behavior occurred across contexts, including those in which freezing is not considered a typical response. We interpret this out-of-context expression of freezing behavior as maladaptive and note its similarity to symptoms of excessive worrying and fear generalization in humans, which are key diagnostic features of ADs (Dunsmoor & Paz, 2015; Dymond et al., 2015; Laufer et al., 2016).

Establishing the long-term efficacy of DREADD-mediated neuronal modulation is important for the development of DREADD technology as a therapeutic approach for human brain disorders, including refractory neuropsychiatric illnesses, which are chronic, disabling, and difficult to treat. While most chemogenetic studies have examined the short-term application of DREADD activation (spanning days to months) in NHPs and rodents, the effectiveness of DREADDs to modulate behavior over a prolonged periods of time (years) remains less certain. A small number of reports suggest that DREADDs remain functional for up to one year in macaques, as supported by [^11^C]clozapine PET imaging (Nagai et al., 2016), resting-state functional connectivity MRI (Grayson et al., 2016), and behavioral measures (Upright & Baxter, 2020). To further address this question, nearly two years following viral vector surgery, we confirmed that hM3Dq expression could be visualized with [^11^C]DCZ PET imaging and that hM3Dq activation could continue to modulate behavior. DREADD activation at this timepoint with either low dose clozapine or DCZ continued to impact anxiety-related behaviors, including increased freezing across contexts. This suggests that hM3Dq receptors not only remain functional at this timepoint, but also that reproducible behavioral changes can be obtained with repeated acute activation of amygdala neurons in our testing paradigm. While we cannot comment on whether levels of DREADD expression changed over time in our study sample, it will be important for future chemogenetic experiments that employ repeated and/or chronic DREADD activation strategies to investigate the degree to which receptor desensitization, as well as changes in neural plasticity, may or may not occur, as these factors could impact behavioral outcomes (Claes et al., 2022).

Similar to our previous study using hM4Di DREADDs (Roseboom et al., 2021), histological analyses of tissue from hM3Dq subjects revealed that DREADD expression occurred most prominently in the BLA. Based on this, and the involvement of the BLA in mediating fear-related behavior, we assume that the hM3Dq-induced changes in anxiety-related behaviors were mediated by BLA neuronal activation. However, because some DREADD expression was found in other amygdala nuclei, as well as in regions outside of the amygdala, it is possible that these regions also contributed to the observed behavioral responses. Within the Ce, which receives input from the BLA and projects to various downstream subcortical regions that mediate fear and anxiety-related responses (Fox, Oler, Tromp, et al., 2015), it was determined that an average of 6% of Ce neurons expressed hM3Dq-HA. This expression level is marginally greater than what we previously observed when using AAV5 in NHPs to express the inhibitory hM4Di DREADD (Roseboom et al., 2021). Previous studies in our laboratory using neurotoxic lesions in NHPs demonstrated a mechanistic role for the Ce in mediating adaptive anxiety-responses (Kalin et al., 2004), and thus, activation of these neurons would be expected to impact anxiety-related behaviors. We also observed sparse levels of DREADD expression in other regions of the anxiety-related neural circuit. These regions include insular cortex, anterior hippocampus, pregenual ACC (Area 25), and regions of orbitofrontal cortex (OPro) (Aggleton, 1986; Aggleton et al., 2015; Barbas et al., 2011; Ghashghaei et al., 2007; Höistad & Barbas, 2008; Kelly et al., 2021; McDonald, 1998; Saunders et al., 1988; Timbie & Barbas, 2014; Wang & Barbas, 2018; Zikopoulos et al., 2017). It is likely that expression occurred in these regions via retrograde and anterograde transport of the AAV5 vector and/or its gene product (Burger et al., 2004; Emborg et al., 2014; Markakis et al., 2010; Paterna et al., 2004; Samaranch et al., 2017). Interestingly, unlike what was reported with the hM4Di-HA construct in Roseboom et al. 2021, relatively little HA immunoreactivity was observed in the bed nucleus of the stria terminalis and sublenticular extended amygdala. These differences in expression pattern might be accounted for by potential differences in trafficking of DREADDs or in titer of the constructs used in our studies (Emborg et al., 2014; Nathanson et al., 2009). It is worth noting that, while substantial correspondence between *in vivo* [^11^C]DCZ binding and *ex vivo* immunohistochemical detection of hM3Dq-HA in the amygdala was observed—even revealing differences in anterior-posterior distribution between subjects—the [^11^C]DCZ PET imaging results were limited in detecting only high levels of DREADD expression in locally targeted neurons, and not lower level expression found at the aforementioned projection sites. The development and use of more specific DREADD radiotracers, such as [^18^F]JHU37107 (Bonaventura et al., 2019), will be advantageous for visualization of DREADD-expressing regions and long-range projection sites.

In summary, this study provides novel data demonstrating that chemogenetic activation of the NHP amygdala results in heightened anxiety responses. These responses are characterized by increased freezing behavior not only in its expected context, but also in contexts where freezing is considered maladaptive. This inappropriate expression of anxiety-related behaviors is similar to the pervasive experience of anxiety and associated avoidance behaviors in humans with ADs. Our data suggest that overactivity of the NHP amygdala is sufficient to produce pathological anxiety and that this paradigm can be a useful model for exploring the pathophysiology of human ADs. We also confirm the utility of *in vivo* PET imaging to validate DREADD expression in living subjects and demonstrate the long-term ability of DREADD activation to impact behavior. Taken together, these findings in NHPs support the development of translational efforts aimed at chemogenetically modulating the function of specific circuits for treating human brain and neuropsychiatric disorders.

## Acknowledgements

We thank the personnel of the Harlow Center for Biological Psychology, the HealthEmotions Research Institute, the Waisman Laboratory for Brain Imaging and Behavior, the Wisconsin National Primate Research Center, the Wisconsin Institutes for Medical Research, K. Brunner, W. Block, M. Emborg, E. Fekete, D. French, and C. Boettcher. Confocal microscopy was performed at University of Wisconsin-Madison (UW) Biochemistry Optical Core, which was established with support from the UW Department of Biochemistry Endowment. This work was supported by NIH grant R01-MH046729 to NHK, as well as grants to the Wisconsin National Primate Center Research Center (P51-OD011106 and P51-RR000167), the Emory National Primate Research Center (P51-OD011132), the Waisman Center (P30-HD003352) and the NIDA intramural research program (ZIA000069). The graphical abstract was created with BioRender.com.

## Author contributions

N.H.K., P.H.R., J.A.O., and S.A.L.M. conceptualized the study. N.H.K. oversaw the study. M.K.R., V.R.E., M.E.O., J.A.O., and S.A.L.M. performed the surgeries. PHR performed the plasma endocrine assays. M.A.B. and M.M. determined plasma clozapine and DCZ levels. M.K.R. and V.R.E. performed the behavioral data collection. N.A., S.A.L.M., J.A.O., M.M.K., and P.H.R. analyzed the behavioral data. S.A.L.M. performed the immunohistochemistry, confocal microscopy, the stereological analyses. A.G. and X.H. performed the electron microscopy and related analyses. M.E.O performed the semi-automated 3D segmentation of infusate delivery regions. A.H.D. and B.T.C. synthesized the PET ligand [^11^C]DCZ and aided in analysis of PET imaging data. J.A.O. created the [^11^C]DCZ group-difference and individual-subject images. S.A.L.M, J.A.O., P.H.R., and N.H.K. wrote the paper. All authors reviewed and provided feedback on the paper.

