## Supplemental Information for "DREADD-mediated amygdala activation is sufficient to induce anxiety-like responses in young nonhuman primates"

Supplemental Contents

### Subject Experimental Details

**Supplemental Table 1. Summary of experiments performed in a given subject.** Subjects E1-E5 are referred to in the text as hM3Dq subjects, while subjects C1-C5 are referred to as unoperated cage-mate controls.

| **Subject ID** | **Surgery** | **ACTH and cortisol** | **Immunohistochemistry** | **Electron microscopy** | **Experiment 1: HIP testing - Clozapine** | **[^11^C]DCZ-µPET** | **Experiment 2: HIP testing – Clozapine and DCZ** |
| --- | --- | --- | --- | --- | --- | --- | --- |
| E1 | ✔ | ✔ | ✔ | - | ✔ | ✔ | - |
| E2 | ✔ | ✔ | - | - | ✔ | - | ✔ |
| E3 | ✔ | ✔ | ✔ | ✔ | ✔ | ✔ | ✔ |
| E4 | ✔ | ✔ | - | - | ✔ | - | ✔ |
| E5 | ✔ | ✔ | ✔ | - | ✔ | ✔ | ✔ |
| C1 | - | ✔ | - | - | ✔ | ✔ | - |
| C2 | - | ✔ | - | - | ✔ | ✔ | ✔ |
| C3 | - | ✔ | - | - | ✔ | - | ✔ |
| C4 | - | ✔ | - | - | ✔ | - | ✔ |
| C5 | - | ✔ | - | - | ✔ | - | ✔ |

### Assessing neuroendocrine effects of clozapine

To examine the effects of clozapine dose (0.03 and 0.1 mg/kg) on HPA axis function, plasma levels of cortisol and ACTH were assessed from samples collected immediately after the NEC context of pre-surgical/baseline testing (at approximately the same time of day; range 8:40 AM – 10:50 AM). To obtain plasma, blood was collected in EDTA tubes and immediately centrifuged at 1,900 x g for 10 min at 4°C and the supernatant collected.

***ACTH assay:*** Plasma samples were assayed for ACTH using the MD Biosciences (Oakdale, MN) enzyme-linked immunosorbent assay (ELISA) following the manufacturer’s instructions. Samples were assayed in duplicate. The inter-assay CV%s were calculated for a low and a high control sample. The low control had an average value of 41.7 ± 0.8 pg/ml and a CV% of 9.0 and the high control had an average value of 249.4 ± 4.2 pg/ml and a CV% of 7.8. The limit of detection defined by the lowest standard was 5 pg/ml.

***Cortisol assay:*** Plasma samples were assayed for cortisol in duplicate using the MP Biomedicals (Solon, OH) Immuchem coated tube radioimmunoassay. The intra-assay CV% was 4.9 and the inter-assay CV% was 9.8. The detection limit defined by the lowest standard was 1 µg/dL.

***Analysis:*** Statistical testing was performed and graphical representation of the data were prepared using Prism v. 9.3.1 (GraphPad Software, San Diego, CA, USA).

***Results:*** Clozapine at either dose used in this study (0.03 and 0.1 mg/kg) significantly decreased stress-induced plasma ACTH (F_2,18_ = 6.16; p < 0.01), replicating the results in our previous DREADD study (Roseboom et al., 2021). In this sample, however, cortisol concentrations were not significantly affected by either dose of clozapine (F_2,18_ = 1.38; p = 0.278); see Supplemental Figure 1.

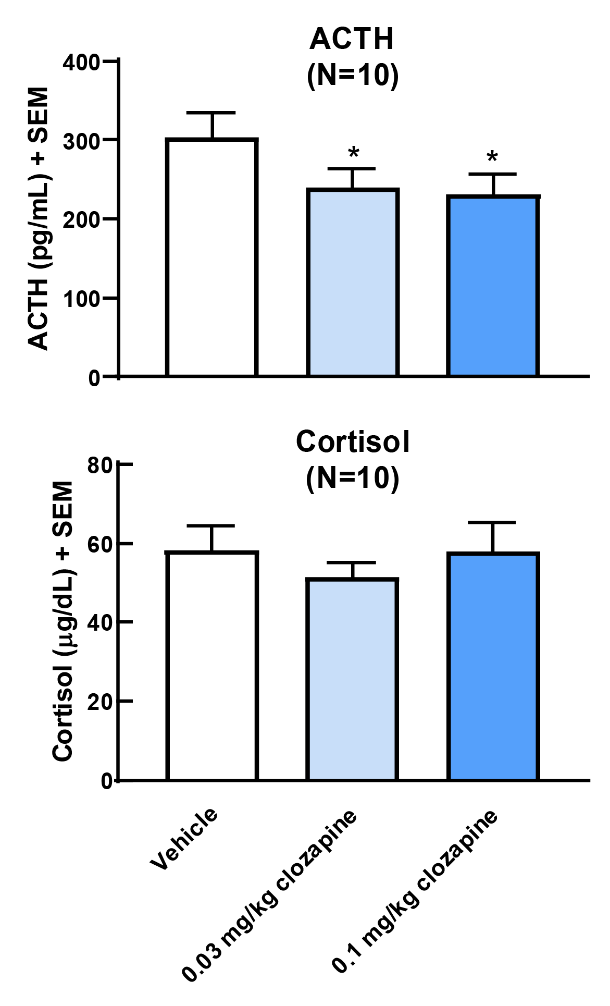
**Supplemental Figure 1. Effects of 0.03 mg/kg and 0.1 mg/kg clozapine on plasma ACTH and cortisol levels in animals selected for this study (n = 10).** ACTH (upper) and cortisol (lower) were assessed following administration of vehicle, 0.03, and 0.1 mg/ kg clozapine to the same animals (n = 10). ACTH was significantly reduced by both doses, whereas no significant reductions in cortisol were detected. (*p < 0.05).

### Semi-automated 3D segmentation of infusate delivery regions

Infusate delivery regions were segmented from iMRI volumes using ITK-SNAP’s 3D segmentation tool operating in threshold mode. Each post-infusion volume was loaded into ITK-SNAP (Supplemental Figure 2A) and examined to empirically determine intensity threshold values such that the thresholded image (Supplemental Figure 2B) would include the gadolinium-enhanced voxels and reject nearby tissue outside the infusate delivery region. The segmentation contour was initialized by placing a small sphere inside the infusate signal (Supplemental Figure 2B). The contour was then allowed to evolve, expanding to include a few additional voxels after 5 iterations (Supplemental Figure 2C) and stabilizing within 120 iterations (Supplemental Figure 2D). All ten infusions in this study were processed in the same way, taking approximately 100 iterations (about 1 second of computation time) until the contour stabilized. The resulting segmentation from the right hemisphere infusion of subject E1, as well as a 3D render of this segmented volume, are shown in Supplemental Figure 2E-H.

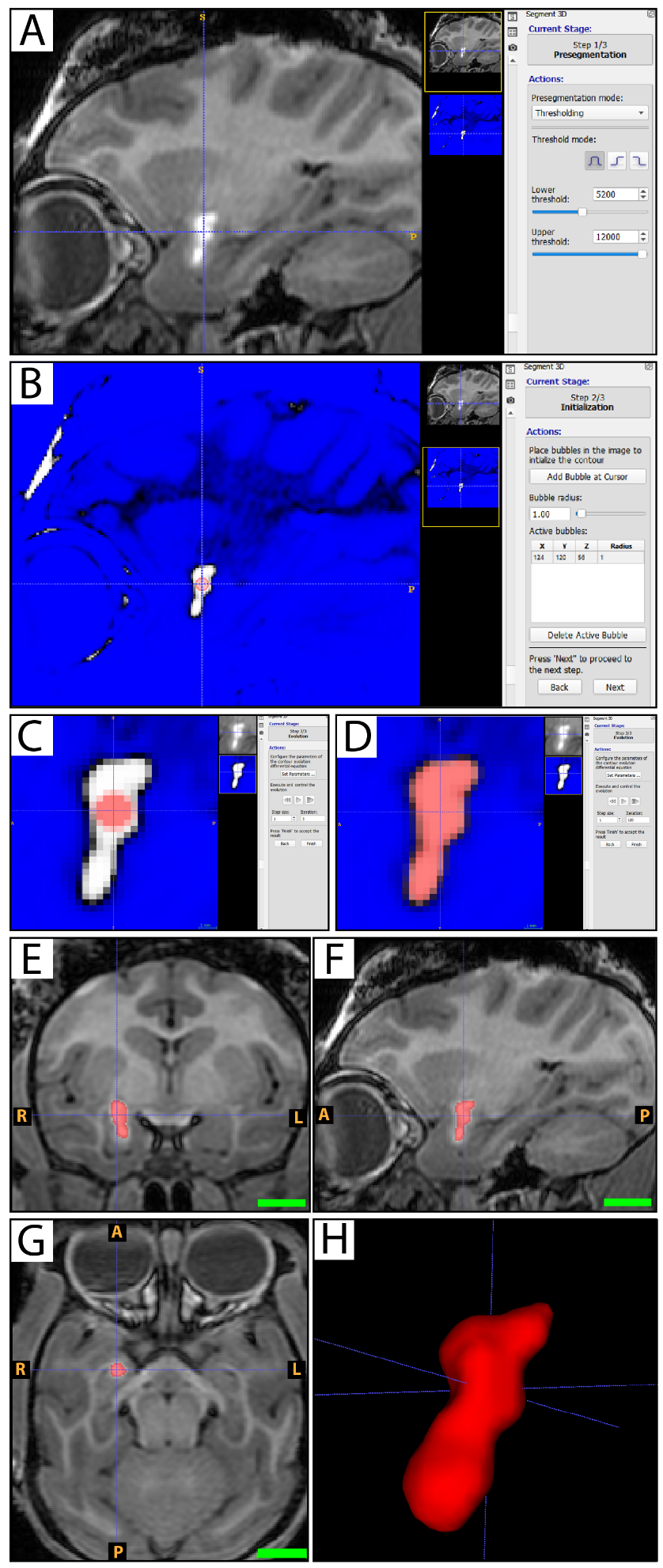
**Supplemental Figure 2. Example of semi-automated 3D infusion segmentation from iMRI volume in subject E1. A)** Sagittal view of post-infusion iMRI showing infusate. **B)** The same image after thresholding, with a blue overlay on excluded voxels. Segmentation contour was initialized with a small sphere (in red). **C)** A zoomed-in view showing that, after five iterations, a few additional voxels have been included as the contour expands outward. **D)** After 120 iterations, segmentation contour has stabilized after expanding to fill the region. Orthogonal slices through the infusate delivery region are shown in **(E)** coronal, **(F)** sagittal, and **(G)** axial views of the segmented voxels (red.) A 3D rendered and shaded surface **(H)** further illustrates the shape of the segmented region. Scalebars (green) in panels E-G represent 1 cm.

### Assessment of hM3Dq-HA expression

#### List of Antibodies

**Supplemental Table 2. Antibodies utilized for immunohistochemical and electron microscopy procedures.**

| **Antibody** | **Host** | **Dilution** | **Source** | **Cat #** |
| --- | --- | --- | --- | --- |
| HA (C29F4) | Rabbit | 1:400 | Cell Signaling Technology, Danvers, MA | 3724 |
| NeuN (clone A60) | Mouse | 1:2000 | MilliporeSigma, Darmstadt, Germany | MAB377 |
| NeuN | Guinea Pig | 1:2000 | Merck KGaA, Darmstadt, Germany | ABN90 |
| GFAP | Rabbit | 1:10000 | DAKO, Agilent, Santa Clara, CA | Z0334 |
| GFAP | Mouse | 1:1000 | Cell Signaling Technology, Danvers, MA | 3670 |
| anti-rabbit HRP | Goat | 1:250 | Vector Laboratories, Burlingame, CA | PI-1000 |
| Biotinylated goat anti-rabbit secondary | Goat | 1:200 | Vector Laboratories, Burlingame, CA | BA-1000 |
| Gold-conjugated anti-rabbit (1.4 nm gold particle size) | Goat | 1:100 | Nanoprobes, Stony Brook, NY | 2004 |
| Alexa Fluor 488 conjugated anti-rabbit secondary | Goat | 1:250 | Invitrogen, ThermoFisher Scientific | A11008 |
| Alexa Fluor 488 conjugated anti-mouse secondary | Donkey | 1:250 | Invitrogen, ThermoFisher Scientific | A21202 |
| Alexa Fluor 568 conjugated anti-rabbit secondary | Goat | 1:250 | Invitrogen, ThermoFisher Scientific | A11011 |
| Alexa Fluor 568 conjugated anti-mouse secondary | Donkey | 1:250 | Invitrogen, ThermoFisher Scientific | A10037 |
| Alexa Fluor 647 conjugated anti-guinea pig secondary | Goat | 1:250 | Invitrogen, ThermoFisher Scientific | A21450 |
| Alexa Fluor 647 conjugated anti-mouse secondary | Donkey | 1:250 | Invitrogen, ThermoFisher Scientific | A31571 |

#### Regional Localization of hM3Dq-HA

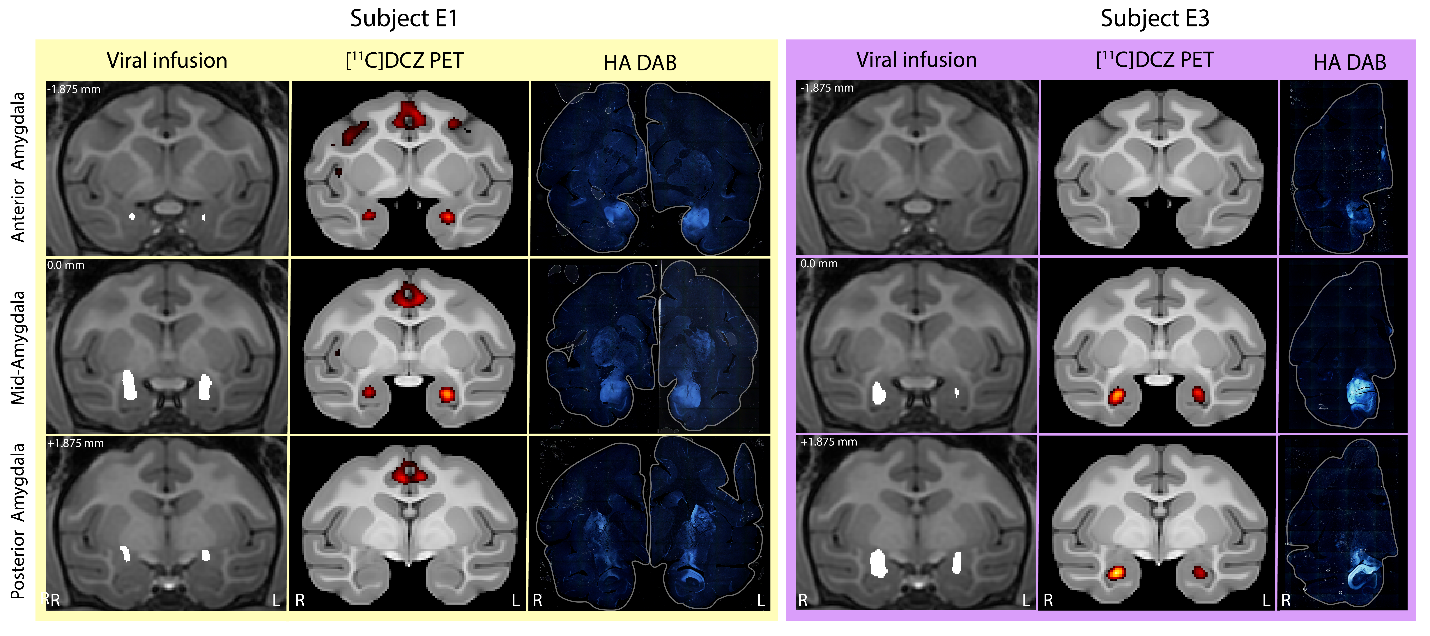

**Supplemental Figure 3. Detection of hM3Dq-HA expression in the NHP amygdala across multimodal measures.** Images showing viral infusion, elevated [^11^C]DCZ PET binding, and immunohistochemical detection of HA-tag are juxtaposed to demonstrate correspondence of these measures within subjects E1 and E3. Viral infusion images show segmented voxels (white) indicative of gadolinium signal, which was used to monitor infusion of AAV5-hSyn-HA-hM3Dq. [^11^C]DCZ PET imaging (thresholded at ≥80% of subject’s peak binding value) demonstrate binding in the amygdala. Immunoperoxidase DAB labeling was used to visualize HA-tag post-mortem, shown in full coronal slice view. HA-DAB is shown only in the right hemisphere for Subject E3, as the left hemisphere was utilized for electron microscopy analyses. Zero (0.0 mm) corresponds to the posterior edge of the anterior commissure. R, right hemisphere; L, left hemisphere.

##### Stereological Assessment of hM3Dq-HA expression in the Ce

**Table 3. Stereological parameters used to assess CeA hM3Dq-HA neuronal expression.**

| Subject | E1 | E3 | E5 |
| --- | --- | --- | --- |
| Hemisphere | L | R | L |
| Section interval (every nth section evaluated) | 10 | 15 | 15 |
| Number of tissue sections evaluated | 5 | 5 | 5 |
| Optical disector height (µm) | 20 | 20 | 20 |
| Sampling % NeuN | 20 | 20 | 20 |
| Sampling % HA+NeuN+ | 100 | 100 | 100 |

**Table 4. Stereological estimates of CeA hM3Dq-HA neuronal expression**.

| Subject | E1 | | | E3 | | | E5 | | |
| --- | --- | --- | --- | --- | --- | --- | --- | --- | --- |
| Region | CeL | CeM | Ce | CeL | CeM | Ce | CeL | CeM | Ce |
| Measured defined mounted thickness (NeuN) in µm | 23.1 | 23.1 | 23.1 | 28.6 | 29.2 | 28.9 | 26.8 | 27.0 | 26.9 |
| Measured defined mounted thickness (HA) in µm | 23.0 | 23.0 | 23.0 | 29.4 | 29.5 | 29.5 | 26.2 | 26.7 | 26.4 |
| Estimated NeuN population using mean section thickness | 51287 | 80641 | 131918 | 89019 | 86649 | 175656 | 99377 | 145340 | 244601 |
| Estimated HA+NeuN+ population using mean section thickness | 2293 | 4923 | 7115 | 8251 | 7778 | 16029 | 4962 | 5919 | 10881 |
| % Transduction Efficiency Estimate | 4.47 | 6.10 | 5.39 | 9.27 | 8.98 | 9.13 | 4.99 | 4.07 | 4.45 |
| NeuN population estimation coefficient of error (Gunderson m=1) | 0.04 | 0.04 | 0.03 | 0.03 | 0.03 | 0.03 | 0.03 | 0.03 | 0.03 |
| HA+NeuN+ population estimation coefficient of error (Gunderson m=1) | 0.09 | 0.06 | 0.05 | 0.05 | 0.05 | 0.04 | 0.07 | 0.06 | 0.04 |

##### Assessing co-labeling of HA-tag, NeuN, and GFAP in the site of viral vector infusion

A small but relatively localized amount of astroglial infiltration and neuronal loss in the amygdala was observed in all hM3Dq-HA subjects examined. Immunofluorescent co-labeling of these areas revealed decreased NeuN and HA immunoreactivity and increased GFAP immunoreactivity, particularly in portions of the anterior dorsal basal nucleus (Supplemental Figure 4A-C). While inflammation and loss of neurons may result from mechanical damage occurring due to insertion of the cannula during iMRI surgery, and/or from the infusion itself, this amount of damage was not observed in subjects from the previous hM4Di-HA study, nor in subjects from other AAV studies conducted in our laboratory using similar techniques. Of the NHP chemogenetic studies published to date, neuroinflammation resulting from viral transduction and/or DREADD expression and activation are generally not reported, though it is unclear if this is because no such effects were observed or if these questions were not pursued. A recent study in rats reported inflammation and neuronal loss following DREADD AAV injection at titers comparable to those used in our study (Goossens et al., 2021). This indicates that DREADD expression, and perhaps DREADD activation, may play some role in inducing neurotoxicity. In the current study, we cannot determine the timepoint at which neuronal loss occurred, or whether this had any effect on the observed behavioral results. However, it should be emphasized that the activation of hM3Dq-HA in the amygdala led to significant, reversible, and reproducible changes in anxiety-related behavior, both in the short term (Experiment 1) and long term (Experiment 2), suggesting that the effects of DREADD-mediated amygdala activation cannot be attributed to the neuronal loss observed. As shown in Supplemental Figure 4D, HA-immunopositive fibers were not colocalized with GFAP, but rather with NeuN-immunopositive somata, indicating that hM3Dq-HA expression was primarily occurring in neurons.

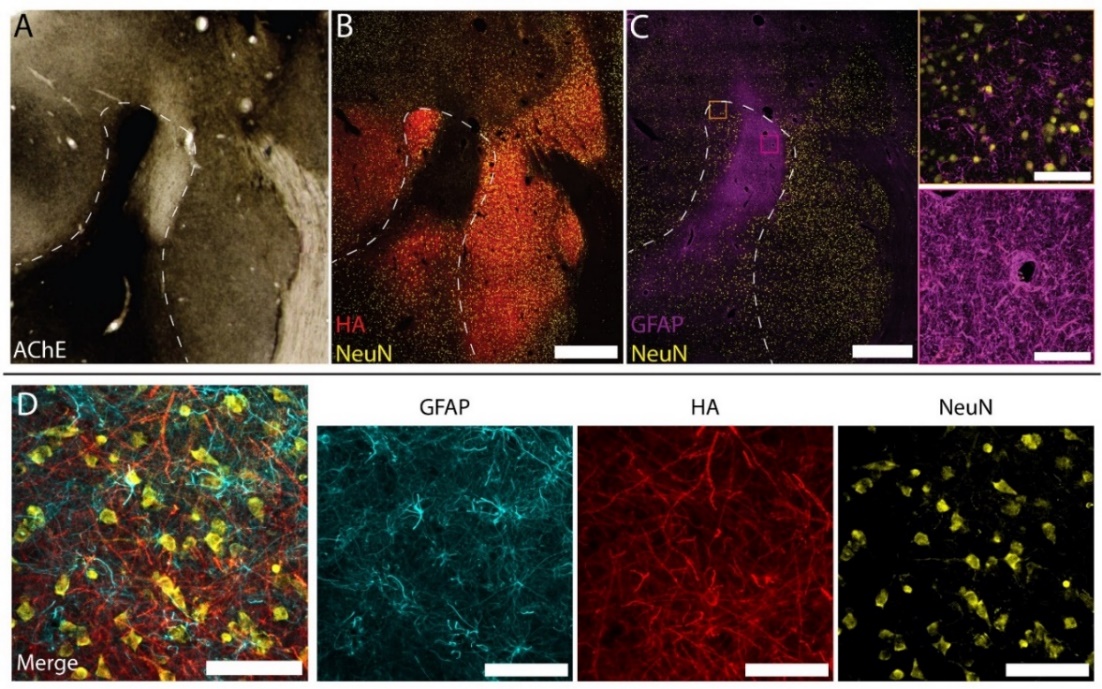

**Supplemental Figure 4. Neuronal loss and astroglial infiltration in the amygdala of subject E5. A)** Dark acetylcholinesterase staining of the basal nucleus is observed, with a void of staining present in the dorsal region. **B)** A nearby section, stained for HA tag and NeuN, shows a lack of HA and NeuN immunoreactivity in this region, and an adjacent section **(C)** reveals an abundance of GFAP, indicative of astroglia. GFAP-immunoreactivity was greatly increased in this region of neuronal loss (pink box), relative to the surrounding region (orange box). **D)** No colocalization of HA-tag and GFAP is observed. Scalebars (white) represent 1000 µm in B and C, or 100 µm in subsets of panel C and in panel D. The basal nucleus is depicted with a white dashed line in panels A-C.

#### Cellular and subcellular localization of hM3Dq-HA

**Table 5. Number of elements quantified in electron microscopy analysis.**

| Region | No. blocks analyzed | No. elements counted | | | | | |
| --- | --- | --- | --- | --- | --- | --- | --- |
|  |  | Dendrites | Glia | Terminals | Unmyelinated axons | Unidentified | Total |
| Central nucleus (lateral) | 2 | 99 | 4 | 77 | 96 | 7 | 283 |
| Central nucleus (medial) | 2 | 92 | 5 | 78 | 243 | 13 | 431 |
| Basal nucleus (dorsal) | 1 | 90 | 1 | 40 | 63 | 12 | 206 |
| Accessory basal nucleus | 1 | 71 | 0 | 31 | 70 | 22 | 194 |

### Experiment 1: Clozapine-induced hM3Dq-HA activation enhances anxiety-related responses

#### Statistical Results

**Supplemental Table 6. F values (and corresponding p-values) of main effects and interactions in Experiment 1: Alone and NEC Contexts**. Significant F values (p<0.05) are bolded.

|  | *Freezing* | | *Locomotion* | | *Cooing* | |
| --- | --- | --- | --- | --- | --- | --- |
|  | *F* | *p* | *F* | *p* | *F* | *p* |
| Group | 0.188 | 0.675 | 1.296 | 0.285 | 4.446 | 0.066 |
| Pre/Post | 0.423 | 0.516 | **8.765** | 0.003 | 2.480 | 0.116 |
| Treatment | **6.475** | 0.011 | 3.496 | 0.062 | 0.472 | 0.492 |
| Condition | **151.909** | 2.20E-16 | **232.682** | 2.20E-16 | 0.007 | 0.935 |
| Group x Pre/Post | **17.142** | 4.29E-05 | **13.378** | 2.91E-04 | 2.421 | 0.121 |
| Group x Treatment | **6.104** | 0.014 | **14.635** | 1.53E-04 | 0.382 | 0.537 |
| Pre/Post x Treatment | 3.308 | 0.070 | **14.339** | 1.78E-04 | 0.087 | 0.768 |
| Group x Condition | 3.492 | 0.062 | 2.406 | 0.122 | 0.130 | 0.718 |
| Pre/Post x Condition | **4.268** | 0.040 | **4.940** | 0.027 | 0.047 | 0.828 |
| Treatment x Condition | 0.364 | 0.547 | **4.837** | 0.028 | 1.468 | 0.226 |
| Group x Pre/Post x Treatment | **4.218** | 0.041 | **15.247** | 1.12E-04 | 0.051 | 0.822 |
| Group x Pre/Post x Condition | 3.277 | 0.071 | 2.811 | 0.094 | 0.248 | 0.619 |
| Group x Treatment x Condition | 1.671 | 0.197 | 0.102 | 0.749 | 1.933 | 0.165 |
| Pre/Post x Treatment x Condition | **4.151** | 0.042 | 3.723 | 0.054 | 0.148 | 0.700 |
| Group x Pre/Post x Treatment x Condition | 0.201 | 0.654 | 2.507 | 0.114 | 0.043 | 0.836 |

**Supplemental Table 7. F values (and corresponding p-values) of main effects and interactions in Experiment 1: Stare Context**. Significant F values (p<0.05) are bolded.

|  | *Freezing* | | *Locomotion* | | *Cooing* | | *Experimenter Hostility* | | *Bark* | |
| --- | --- | --- | --- | --- | --- | --- | --- | --- | --- | --- |
|  | *F* | *p* | *F* | *p* | *F* | *p* | *F* | *p* | *F* | *p* |
| Group | 0.367 | 0.562 | 1.320 | 0.284 | 4.534 | 0.066 | 0.393 | 0.548 | 0.745 | 0.413 |
| Treatment | **20.384** | 2.59E-05 | 2.660 | 0.108 | 1.827 | 0.181 | 2.039 | 0.158 | 1.574 | 0.214 |
| Group x Treatment | **11.271** | 0.001 | **11.762** | 0.001 | 0.193 | 0.662 | 1.748 | 0.191 | 3.925 | 0.052 |

#### Freezing behavior

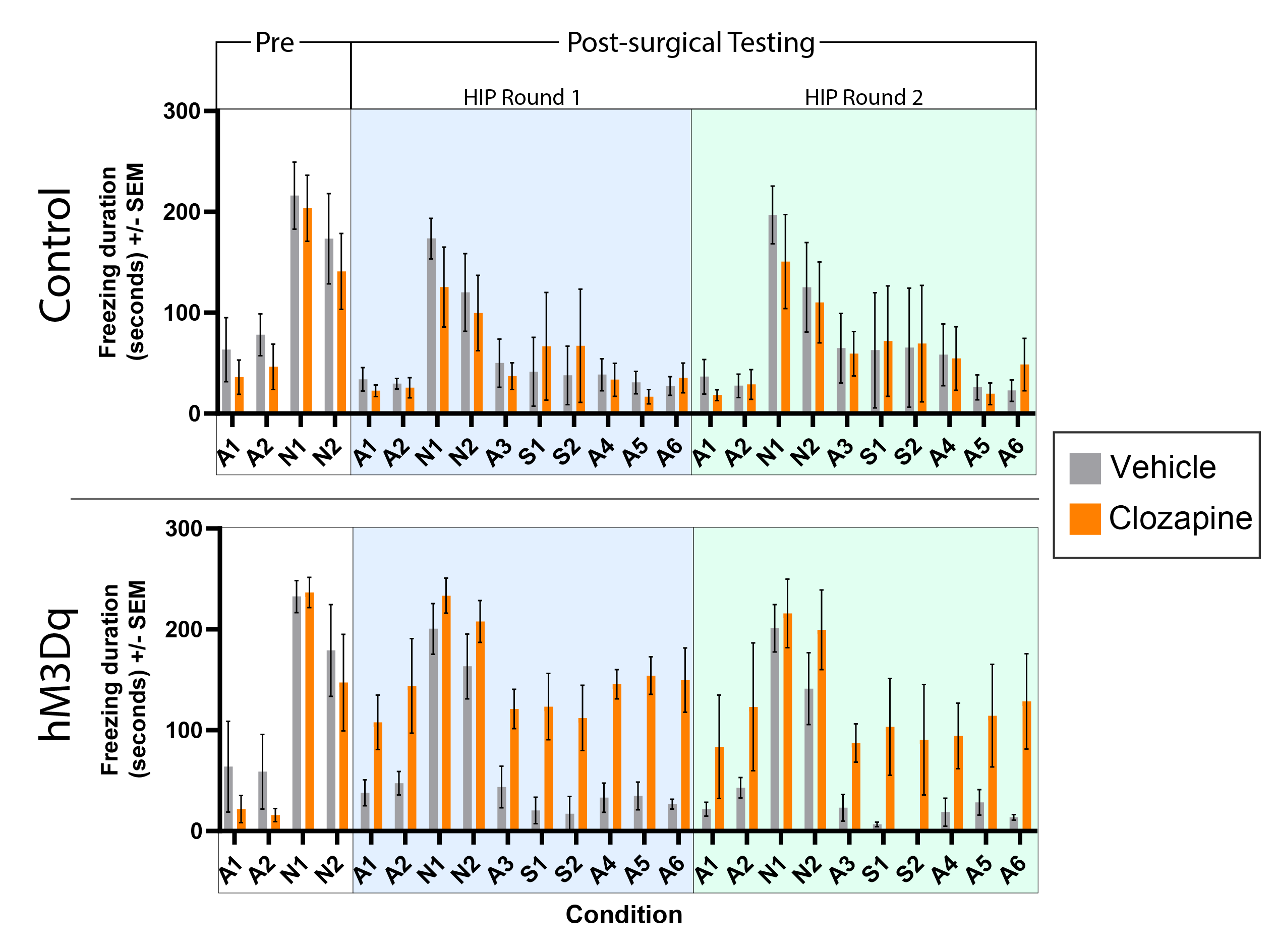

**Supplemental Figure 5. Untransformed freezing duration scores during Experiment 1.** Freezing duration scores are shown for control (top) and hM3Dq subjects (bottom) after vehicle and clozapine administration timepoints throughout Experiment 1. Baseline or “Pre” behavioral testing (conducted prior to surgery/AAV transduction in the hM3Dq group), is shown with a white background, while “Post-surgical” testing is shown with a blue (Round 1) or green (Round 2) background.

#### Locomotion behavior

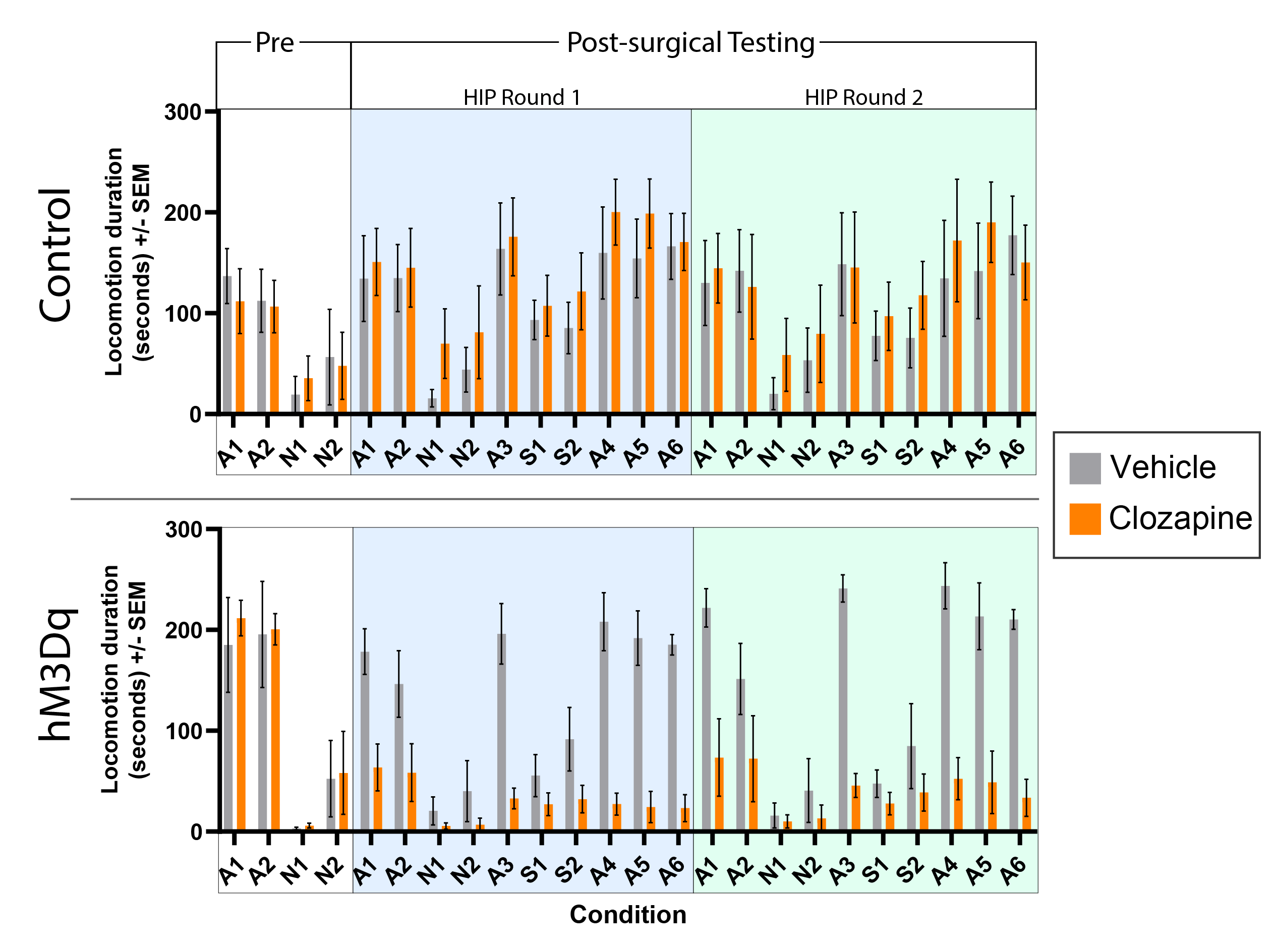

**Supplemental Figure 6. Untransformed locomotion duration scores during Experiment 1.** Locomotion duration scores are shown for control (top) and hM3Dq subjects (bottom) after vehicle and clozapine administration timepoints throughout Experiment 1. Baseline or “Pre” behavioral testing (conducted prior to surgery/AAV transduction in the hM3Dq group), is shown with a white background, while “Post-surgical” testing is shown with a blue (Round 1) or green (Round 2) background.

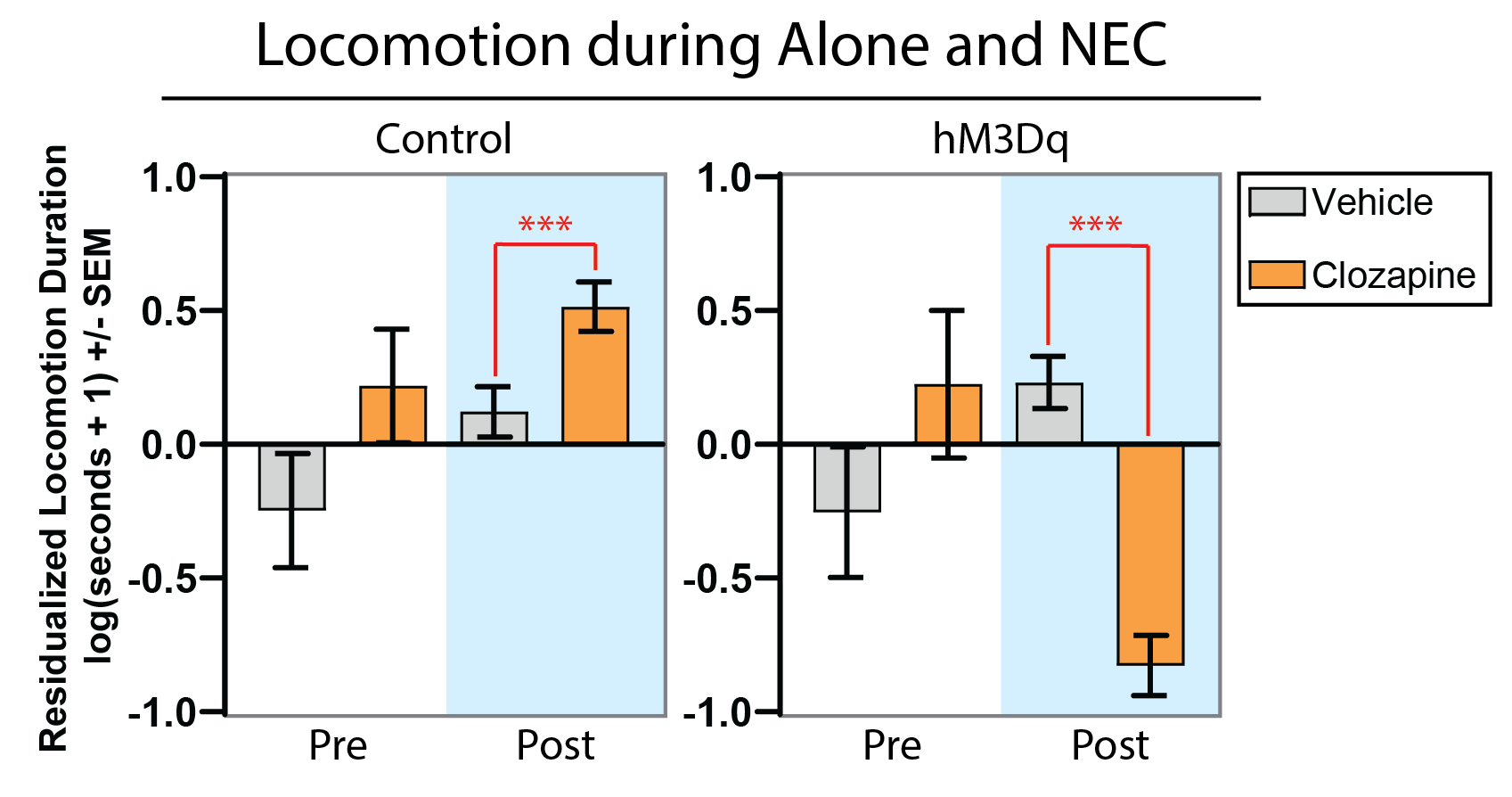

**Supplemental Figure 7. hM3Dq-mediated amygdala activation decreases locomotion behavior during the Alone and NEC conditions**. Graphs show mean log-transformed locomotion duration (residualized for plasma clozapine concentrations) across the Alone and NEC conditions of the HIP. Data following vehicle or clozapine administration are shown for both the control and hM3Dq groups during pre-surgical (white background) and post-surgical (blue background) testing. A significant Group x Treatment x Pre/Post interaction was observed (F_1,376_ = 15.247, p < 0.001). ***p<0.001

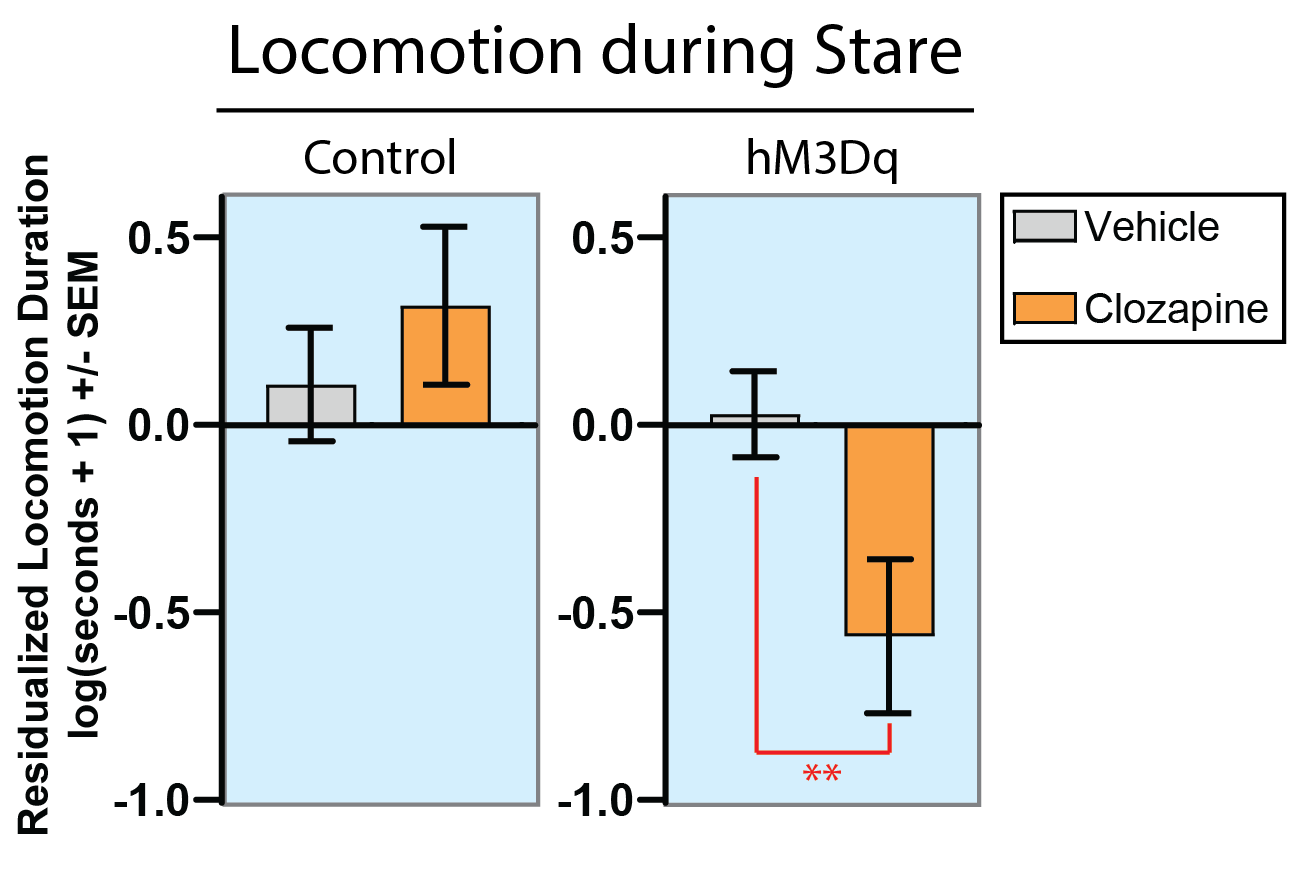

**Supplemental Figure 8. hM3Dq-mediated amygdala activation decreases locomotion behavior during the Stare condition**. Mean log-transformed locomotion duration (residualized for plasma clozapine concentrations) following vehicle or clozapine administration is shown for both the control and hM3Dq groups during the Stare condition. A significant Group x Treatment interaction was observed (F_1,68_ = 11.762, p = 0.001). (**p<0.01)

### Experiment 2: Anxiety-related responses continue to be affected by DREADD-mediated amygdala activation nearly two years following viral vector infection

#### Statistical Results

**Supplemental Table 8. F values (and corresponding p-values) of main effects and interactions in Experiment 2: Clozapine-mediated hM3Dq activation**. Significant F values (p<0.05) are bolded.

|  |  | *Freezing* | | | *Locomotion* | | *Cooing* | | *Experimenter Hostility* | | *Bark* | |
| --- | --- | --- | --- | --- | --- | --- | --- | --- | --- | --- | --- | --- |
|  |  | *F* | *p* | *F* | | *p* | *F* | *p* | *F* | *p* | *F* | *p* |
|  | Group | 0.353 | 0.571 | 0.058 | | 0.817 | 1.247 | 0.306 | 1.253 | 0.306 | 1.081 | 0.339 |
|  | Treatment | 3.503 | 0.063 | 0.346 | | 0.557 | 1.356 | 0.246 | 3.165 | 0.089 | 0.909 | 0.351 |
|  | Condition | **96.707** | 2.20E-16 | **103.766** | | 2.20E-16 | **9.081** | 1.94E-04 | - | - | - | - |
|  | Group x Treatment | **13.366** | 3.60E-04 | **16.814** | | 6.91E-05 | **4.930** | 0.028 | 0.341 | 0.565 | **5.538** | 0.028 |
|  | Group x Condition | **13.123** | 5.90E-06 | **4.424** | | 0.014 | 0.050 | 0.951 | - | - | - | - |
|  | Treatment x Condition | **5.854** | 0.004 | 0.100 | | 0.905 | 0.144 | 0.866 | - | - | - | - |
|  | Group x Treatment x Condition | 2.742 | 0.068 | 1.096 | | 0.152 | 0.710 | 0.493 | - | - | - | - |

**Supplemental Table 9. F values (and corresponding p-values) of main effects and interactions in Experiment 2: DCZ-mediated hM3Dq activation**. Significant F values (p<0.05) are bolded.

|  | *Freezing* | | *Locomotion* | | *Cooing* | | *Experimenter Hostility* | | *Bark* | |
| --- | --- | --- | --- | --- | --- | --- | --- | --- | --- | --- |
|  | *F* | *p* | *F* | *p* | *F* | *p* | *F* | *p* | *F* | *p* |
| Group | 0.008 | 0.931 | 0.252 | 0.631 | 0.691 | 0.437 | 2.831 | 0.143 | 4.546 | 0.077 |
| Treatment | 0.121 | 0.729 | 1.715 | 0.192 | 2.079 | 0.152 | 0.000 | 1.000 | 1.679 | 0.208 |
| Condition | **139.465** | 2.20E-16 | **91.113** | 2.20E-16 | **13.806** | 3.32E-06 | - | - | - | - |
| Group x Treatment | **4.756** | 0.031 | **21.241** | 8.94E-06 | 0.013 | 0.911 | 0.690 | 0.415 | **9.313** | 0.006 |
| Group x Condition | **10.769** | 4.42E-05 | 2.228 | 0.111 | 0.274 | 0.761 | - | - | - | - |
| Treatment x Condition | 0.147 | 0.863 | 0.783 | 0.459 | 1.109 | 0.333 | - | - | - | - |
| Group x Treatment x Condition | 1.676 | 0.191 | **3.428** | 0.035 | 0.383 | 0.682 | - | - | - | - |

#### Freezing behavior

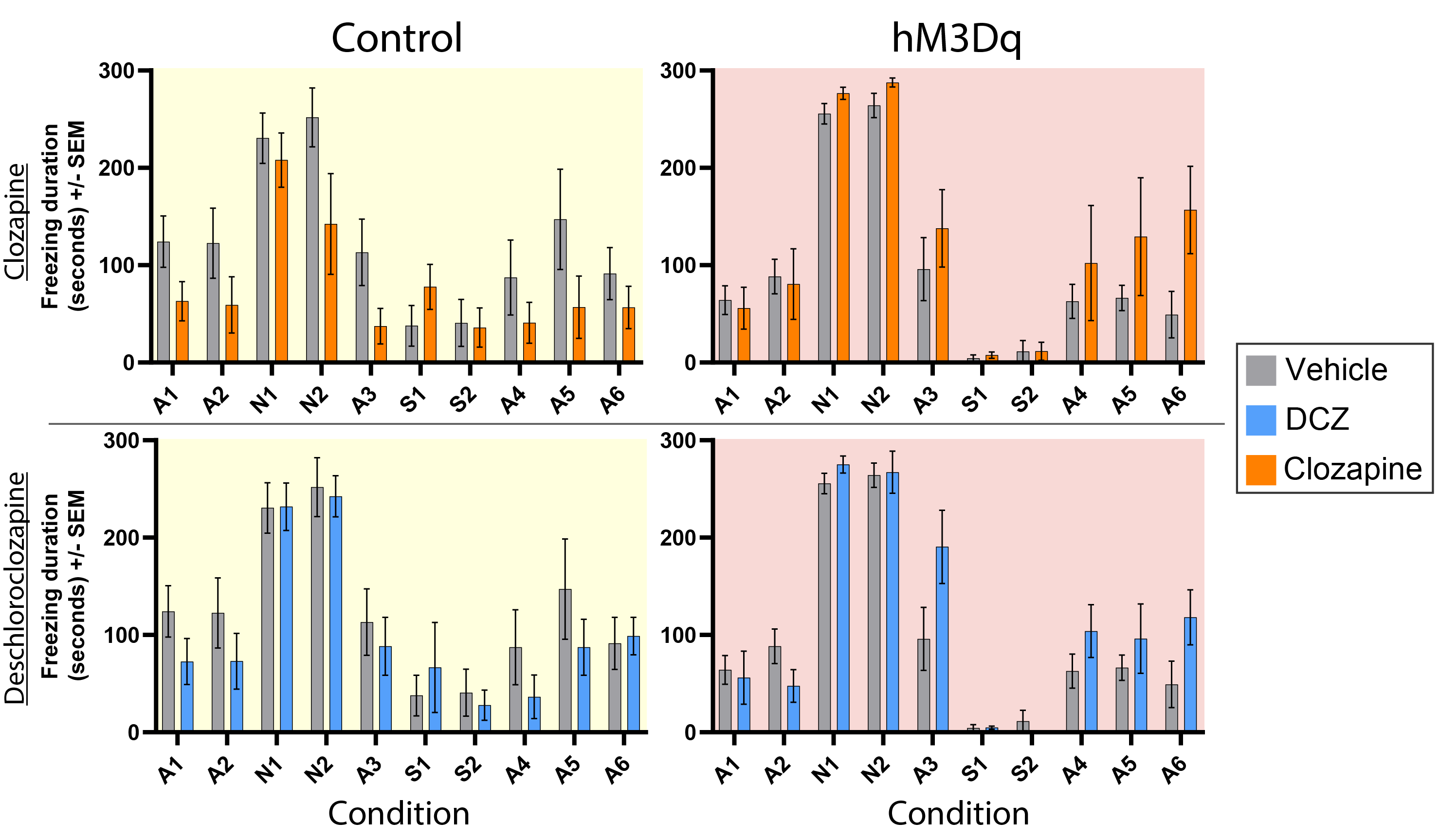

**Supplemental Figure 9. Untransformed freezing durations during Experiment 2.** Freezing duration scores are shown for control (left; yellow background) and hM3Dq subjects (right; red background) after administration of vehicle (same data in upper and lower panels), clozapine (upper panels), or DCZ (lower panels).

#### Locomotion behavior

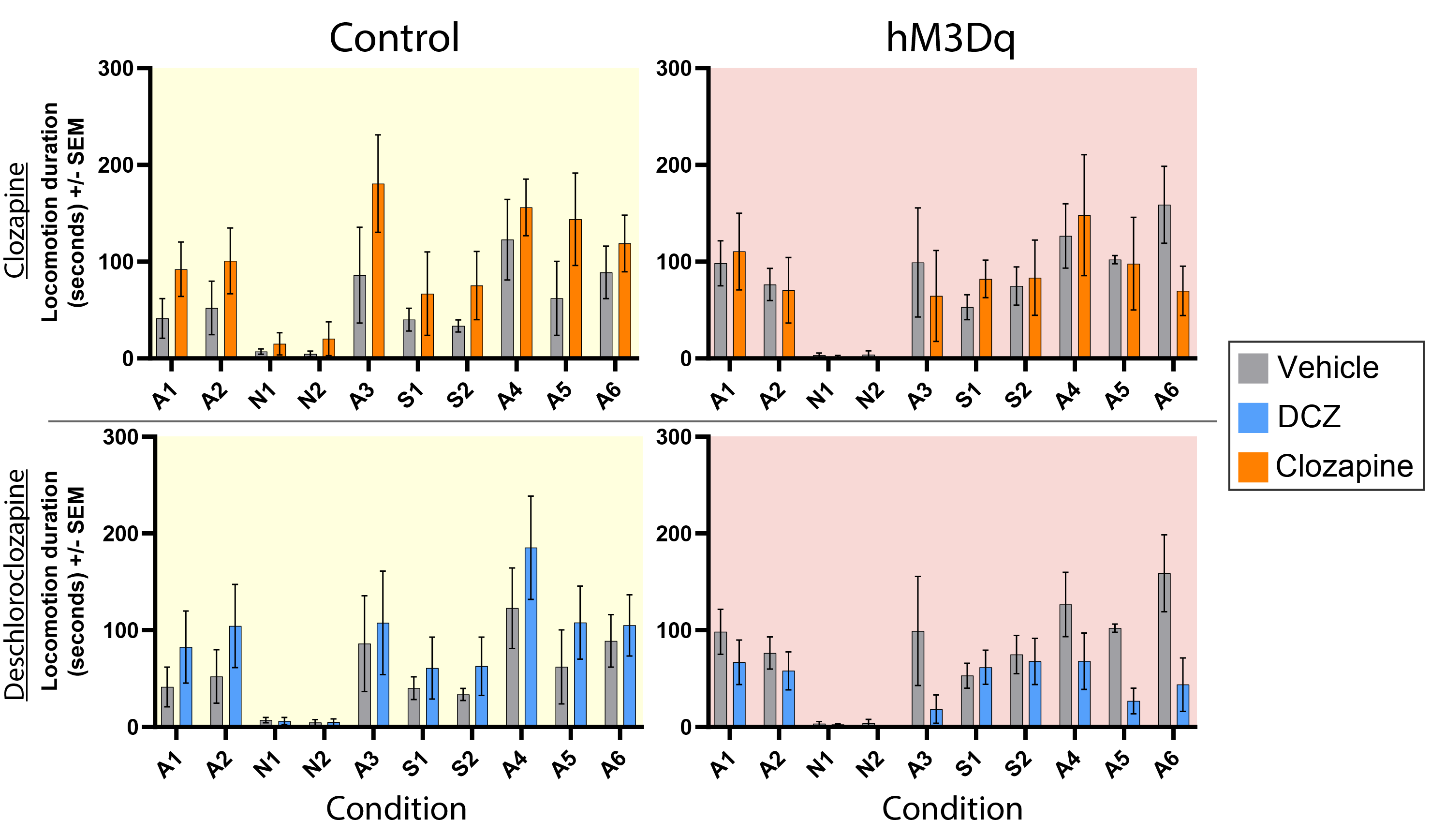

**Supplemental Figure 10. Untransformed locomotion durations during Experiment 2.** Locomotion duration scores are shown for control (left; yellow background) and hM3Dq subjects (right; red background) after administration of vehicle (same data in upper and lower panels), clozapine (upper panels), or DCZ (lower panels).

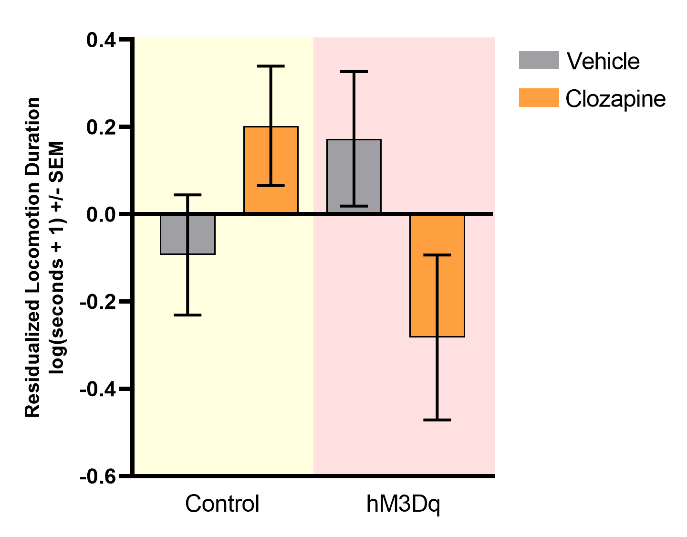
**Supplemental Figure 11. Clozapine-mediated hM3Dq activation continues to induce decreases in locomotion after long-term hM3Dq expression**. Mean log-transformed locomotion duration (residualized for plasma clozapine concentrations) following vehicle or clozapine administration is shown for both the control (yellow background) and hM3Dq (red background) groups. The significant Group x Treatment interaction (F_1,142_ = 16.814, p < 0.001) suggests that the effects of DREADD activation occur across contexts. *Post hoc* comparisons revealed no significant treatment-related differences in either group.

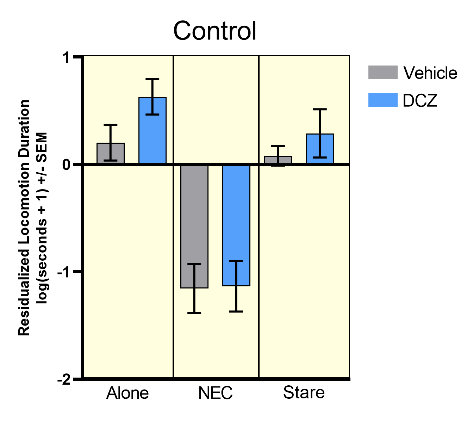

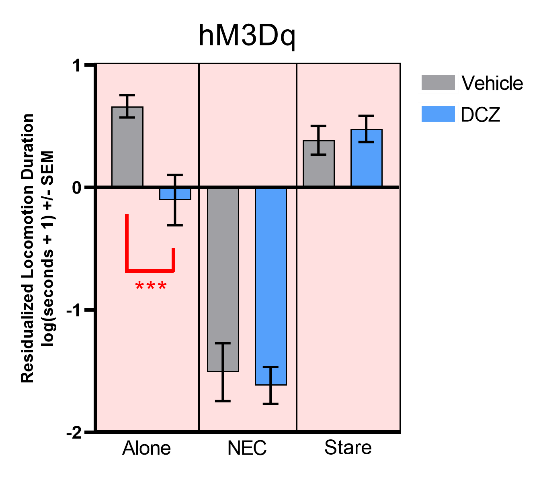

**Supplemental Figure 12. DCZ-mediated hM3Dq activation continues to induce decreases in locomotion after long-term hM3Dq expression, selectively during the Alone condition**. Mean log-transformed locomotion durations (residualized for plasma clozapine concentrations) following vehicle or DCZ administration are shown for both the control (yellow background) and hM3Dq (red background) groups. The significant Group x Treatment x Condition interaction (F_2,142_ = 3.428, p < 0.05), suggests that the effects of DREADD activation occur in a context-specific manner. *Post hoc* comparisons revealed significant treatment-related differences in the hM3Dq group during the Alone condition. (***p<0.001)

#### Coo vocalizations

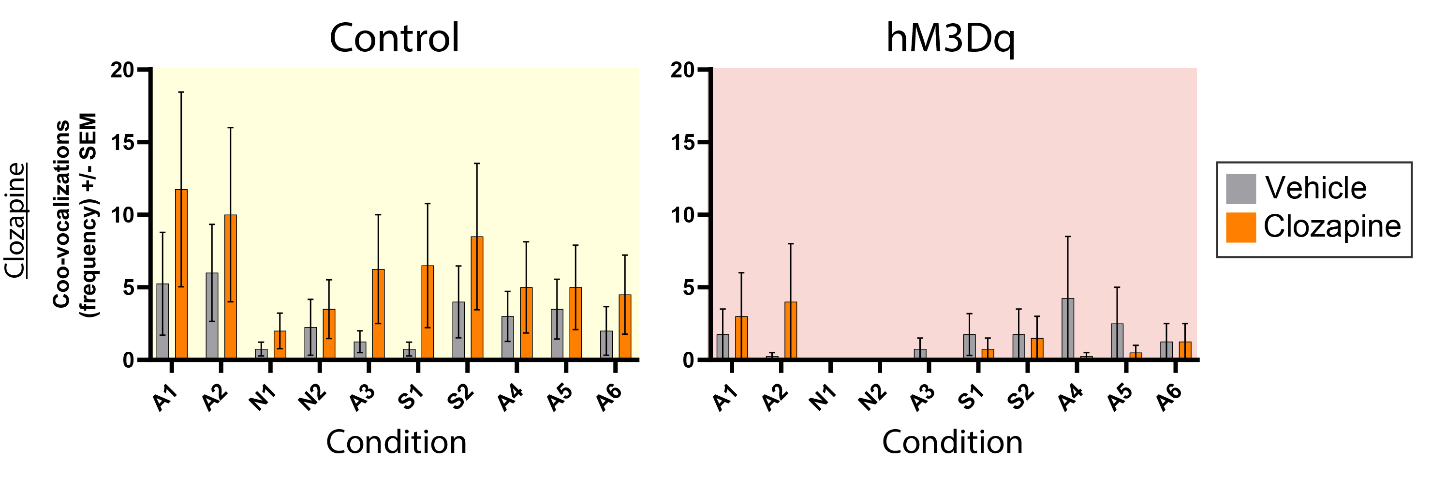

**Supplemental Figure 13. Untransformed frequency of coo-vocalizations during Experiment 2.** Frequency scores of coo-vocalizations are shown for control (left; yellow background) and hM3Dq subjects (right; red background) after administration of vehicle or clozapine.

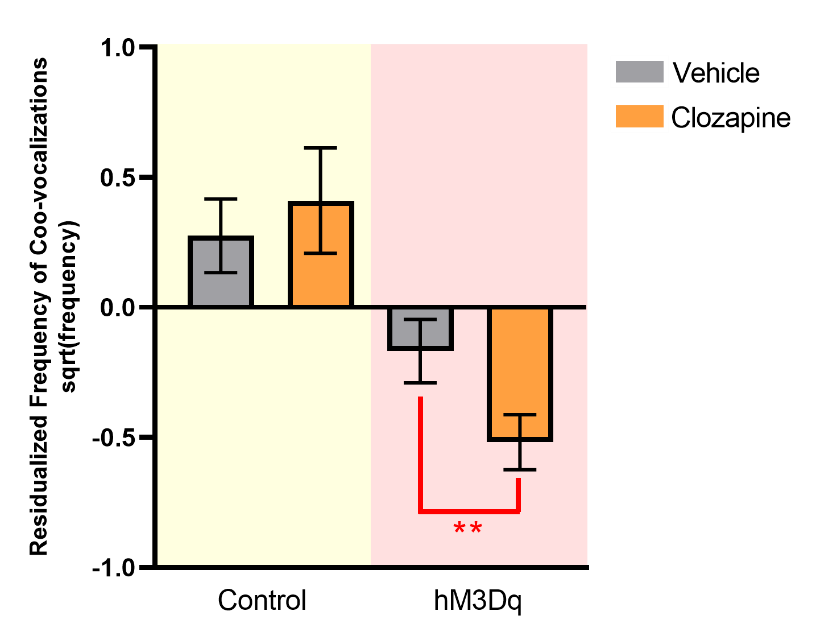
**Supplemental Figure 14. Clozapine-mediated hM3Dq activation decreases coo-vocalizations after long-term hM3Dq expression**. Mean square root-transformed frequency scores (residualized for plasma clozapine concentrations) following vehicle or clozapine administration are shown for both the control (yellow background) and hM3Dq (red background) groups. The significant Group x Treatment interaction (F_1,142_ = 4.930, p < 0.05) suggests that the effects of DREADD activation occur across contexts. (**p<0.01)

#### Bark vocalizations

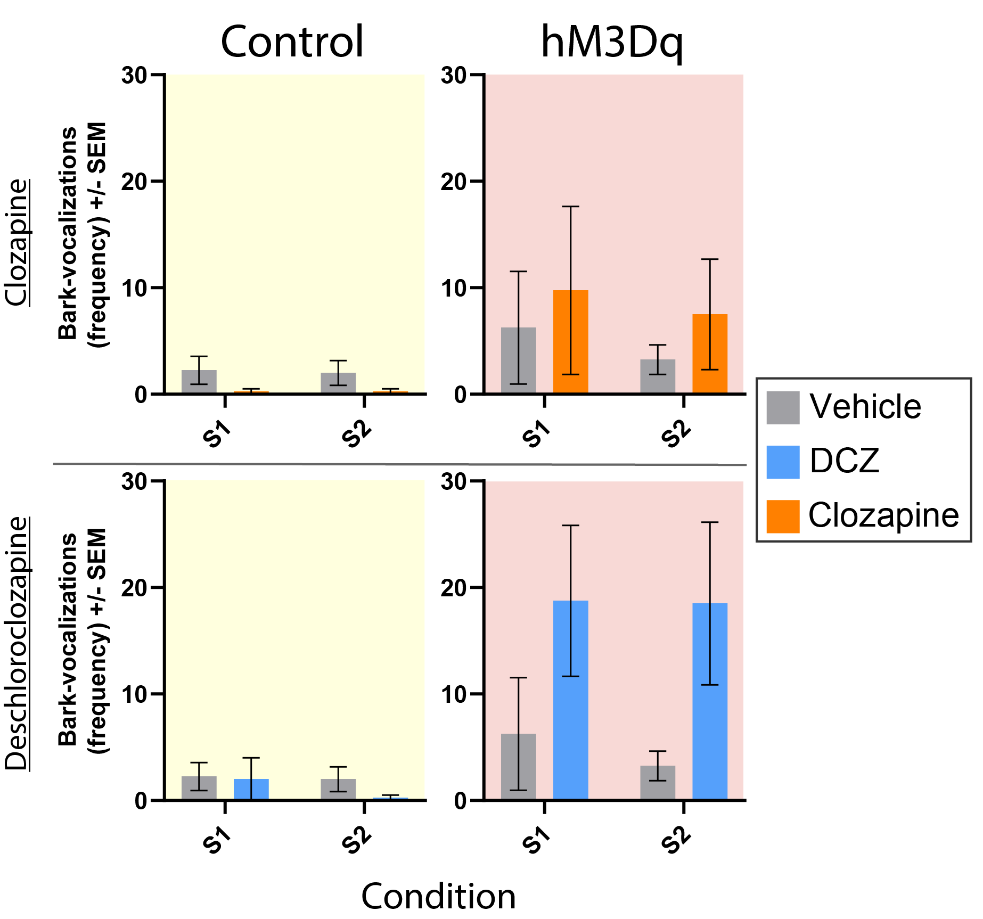
**Supplemental Figure 15. Untransformed frequency of Stare-specific bark-vocalizations during Experiment 2.** Frequency scores of bark-vocalizations are shown for control (left; yellow background) and hM3Dq subjects (right; red background) after administration of vehicle (same data in upper and lower panels), clozapine (upper panels), or DCZ (lower panels).

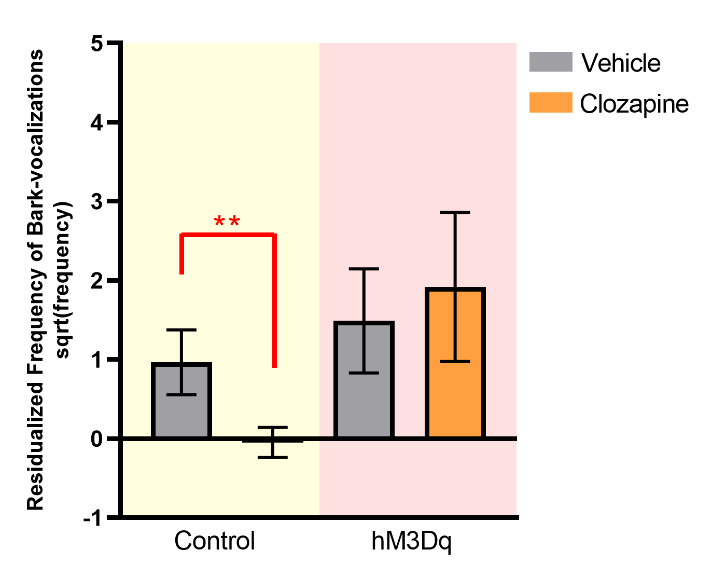
**Supplemental Figure 16. Clozapine reduces Stare-specific bark-vocalizations in control subjects but not in hM3Dq subjects after long-term hM3Dq expression.** Mean square root-transformed frequency scores (residualized for plasma clozapine concentrations) following vehicle or clozapine administration are shown for both the control (yellow background) and hM3Dq (red background) groups. A significant Group x Treatment x Pre/Post interaction was observed (F_1,22_ = 5.538, p < 0.05), such that control subjects barked significantly less on clozapine, relative to vehicle, while no such change was observed in hM3Dq subjects. (**p<0.01)

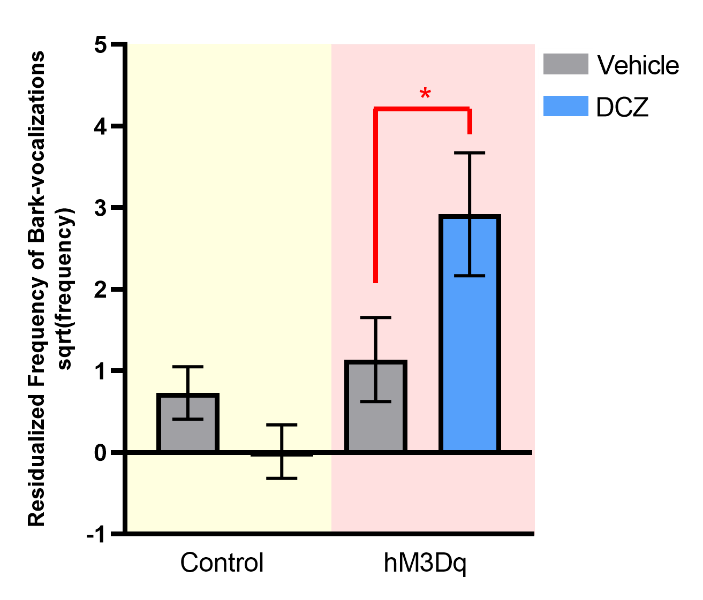
**Supplemental Figure 17. DCZ-mediated hM3Dq activation increases Stare-specific bark-vocalizations after long-term hM3Dq expression.** Mean square root-transformed frequency scores (residualized for plasma clozapine concentrations) following vehicle or clozapine administration are shown for both the control (yellow background) and hM3Dq (red background) groups. A significant Group x Treatment x Pre/Post interaction was observed (F_1,22_ = 9.313, p < 0.01), such that hM3Dq subjects barked significantly more on clozapine, relative to vehicle, while no such change was observed in controls. (*p<0.05)
